# Unraveling Neuronal Identities Using SIMS: A Deep Learning Label Transfer Tool for Single-Cell RNA Sequencing Analysis

**DOI:** 10.1101/2023.02.28.529615

**Authors:** Jesus Gonzalez-Ferrer, Julian Lehrer, Ash O’Farrell, Benedict Paten, Mircea Teodorescu, David Haussler, Vanessa D. Jonsson, Mohammed A. Mostajo-Radji

## Abstract

Large single-cell RNA datasets have contributed to unprecedented biological insight. Often, these take the form of cell atlases and serve as a reference for automating cell labeling of newly sequenced samples. Yet, classification algorithms have lacked the capacity to accurately annotate cells, particularly in complex datasets. Here we present SIMS (Scalable, Interpretable Machine Learning for Single-Cell), an end-to-end data-efficient machine learning pipeline for discrete classification of single-cell data that can be applied to new datasets with minimal coding. We benchmarked SIMS against common single-cell label transfer tools and demonstrated that it performs as well or better than state of the art algorithms. We then use SIMS to classify cells in one of the most complex tissues: the brain. We show that SIMS classifies cells of the adult cerebral cortex and hippocampus at a remarkably high accuracy. This accuracy is maintained in trans-sample label transfers of the adult human cerebral cortex. We then apply SIMS to classify cells in the developing brain and demonstrate a high level of accuracy at predicting neuronal sub-types, even in periods of fate refinement, shedding light on genetic changes affecting specific cell types across development. Finally, we apply SIMS to single cell datasets of cortical organoids to predict cell identities and unveil genetic variations between cell lines. SIMS identifies cell-line differences and misannotated cell lineages in human cortical organoids derived from different pluripotent stem cell lines. When cell types are obscured by stress signals, label transfer from primary tissue improves the accuracy of cortical organoid annotations, serving as a reliable ground truth. Altogether, we show that SIMS is a versatile and robust tool for cell-type classification from single-cell datasets.

## 1. Introduction

Next-generation sequencing systems have allowed for large scale collection of transcriptomic data at the resolution of individual cells. Within this data lies variability allowing us to uncover cell-specific features, such as cell type, state, regulatory networks, as well as infer trajectories of cell differentiation and specification [1, 2]. These properties are crucial to understand biological processes in healthy and diseased tissue. In addition, these properties better inform the development of *in vitro* models, which are often benchmarked against cell atlases of primary tissue [1].

The lowering costs of sequencing, coupled with several barcoding strate-gies, have allowed single-cell datasets and atlases to scale with respect to cell and sample numbers, as well as data modalities [3]. Yet, despite the increas-ing size and complexity of datasets, the most popular pipelines for single cell analysis are based on dimensionality reduction and unsupervised clustering followed by manual interpretation and annotation of each cell cluster [4]. This requires a high level of expertise in understanding the most appropriate cell markers for a given tissue, a major barrier to newcomers to a field. For highly heterogeneous tissues such as the brain, where a consensus in cell type nomenclature remains challenging [5], manual cell annotation can introduce additional errors.

Errors in cell annotation may be driven by the following common as-sumptions: 1) That marker genes are uniformly highly expressed, which is not always the case [6, 7]. For instance, while OPALIN and HAPLN2 are considered markers of oligodendrocytes in the brain, their expression is low or undetectable in a large subset of oligodendrocytes at the single cell level [8]. Indeed, high levels of HAPLN2 have been proposed as a landmark of Parkinson’s Disease [9]. 2) That cell-type marker gene expression is constant throughout development, such that a gene that specifically labels a population of cells at one age would label the same population at a different age. For example, while it is known that PVALB positive cortical interneurons are born during embryonic development [10], the expression of this gene is not seen until well after birth [11]. Notably, recent studies have shown that a subset of PVALB interneurons may never express the PVALB gene [12]. 3) That gene markers discovered in one species apply to others. In several tissues, including the brain, there are major species-specific differences. For example, HCN1 is a key marker of cortical layer 5 sub-cerebral projection neurons in the mouse, but highly expressed in projection neurons of all cortical layers in humans [13, 14]. In summary, manual annotation of every new dataset based on standard marker genes can lead to compounding error propagation and inconsistent single cell atlases, potentially reducing their utility.

The development of software to automate single cell analysis has become an important and popular research topic [4, 15, 16, 17]. However, the accuracy of these automated classifiers often degrades as the number of cell types increase, and the number of samples per label becomes small [18].The distribution of cell types is often asymmetric, with a majority class dominating a high percentage of cells. Additionally, technical variability between experiments can make robust classification between multiple tissue samples difficult. There have been efforts to apply statistical modeling to this problem [19, 20], but the high-dimensional nature of transcriptomic data makes analysis statistically and computationally intractable [21]. These conditions make applying classical models such as support vector machines difficult and ineffective[22]. In response, generative neural networks have become a popular framework due to their robustness to technical variability within data, scalability, and ability to capture biological variation in the latent representation of the inputs [23, 24, 25]. These include deep learning models based on variational inference [26, 27], adversarial networks [28] and attention transformers [25]. Early deep learning models exhibit a lack of interpretability due to their “black box” architecture[18]. However, explainable artificial intelligence (XAI) research aims to understand model decision-making by assigning weight values to the genes based on their influence on cell type predictions. Despite this, some deep learning approaches display inherent biases favoring multivariate gene selection that impedes straightforward data interpretation [25, 29]. Additionally, the computational demands of certain deep learning systems may preclude adoption by smaller research groups lacking access to high-performance computing infrastructure. Ongoing work seeks to enhance model interpretability and efficiency to enable broader utilization across the biological sciences[25, 28].

Here we present SIMS (Scalable, Interpretable Machine Learning for Single-Cell), a new framework based on the model architecture found in TabNet [30]. SIMS is implemented in Pytorch Lightning [31], which allows SIMS to be low code and easy to use. We take advantage of the fact that TabNet uses a sequential self attention mechanism, which allows for interpretability of tabular data [30]. Importantly, TabNet does not require any feature preprocessing and has built-in interpretability which visualizes the contribution of each feature to the model [30]. Given these properties, SIMS is an ideal tool to classify RNA sequencing data. We show that SIMS either outperforms or is on par with state of the art single cell classifiers in complex datasets, such as peripheral blood samples and full body atlases. We apply SIMS to datasets of the mammalian brain and show a high accuracy in adult and developing tissue. We further apply SIMS to data generated from *in vitro* models, such as pluripotent stem cell-derived cortical organoids. Using the SIMS pipeline, we were able to reclassify misslabeled cells through the use of label transfer from annotated primary tissue. We propose SIMS as a new label transfer tool, capable of robust performance with deep annotation and skewed label distributions, high accuracy with small and large datasets, and direct interpretability from the input features.

## 2. Results

### 2.1. Development of a TabNet-based framework for label transfer across single cell RNA datasets

We developed SIMS, a framework for label transfer across single cell RNA datasets that uses TabNet as the classifier component (Supplemental Figure 1) [30]. TabNet is a transformer-based neural network with sparse feature masks that allow for direct prediction interpretability from the input features [30]. To better fit the model for the task of single cell classification we added two innovations: First, we included Temperature Scaling, a post-processing step of the train network that provides the users with a calibrated probability measure for the classification of each cell in the selected cell type [32]. Then, we equipped our pipeline with an automated gene intersection mechanism, allowing the prediction of datasets with a different number of genes than the dataset used for training the model, a common occurrence when different sequencing technologies are used.

In our framework, for each forward pass, batch-normalization is applied. The encoder is several steps (parameterized by the user) of self-attention layers and learned sparse feature masks. The decoder then takes these en-coded features and passes them through a fully-connected layer with batch-normalization and a generalized linear unit activation [33]. Interpretability by sample is then measured as the sum of feature mask weights across all encoding layers.

SIMS can be trained with either one or several preannotated input datasets, allowing for the integration of atlases generated by the same group or by different groups. For accurate training, the user must input an annotated matrix of gene expression in each cell. After training and production of train-ing statistics, the user can input a new unlabeled dataset. Of note, if the training data was normalized ahead of training, the user must normalize the unlabeled data in a similar manner. The model will then predict the cluster assignment for each cell. SIMS will then output the probability of each cell belonging to each cluster, where the probability is more than 0.

SIMS is accessible through a Python API. The development version can be found on our GitHub repository at the following link https://github.com/braingeneers/SIMS. Additionally, a pip package is also available for easy installation https://pypi.org/project/scsims/. SIMS is designed to require minimal input from the users. To train the model, the user has to only input the data file of the training dataset, a file with the labels, and define the class label, the user can also choose to load the dataset into Scanpy as an anndata object (Supplemental Figure 2). This process will save the learned parameters for each training epoch in a new file.

To perform the label transfer on a new dataset the user must import the weights from the trained model. The user will then input the new unlabeled dataset (Supplemental Figure 3).

SIMS takes the cell by gene expression matrix as an input. For newly produced data we recommend an end to end pipeline we have developed within Terra. This pipeline takes raw FASTQ files, runs them through the CellRanger or StarSolo Dockstore workflows [34, 35, 36] (Supplemental Figure 4), outputs an expression matrix in the h5 format and classifies the cell types using a SIMS model trained on the reference dataset of interest. This pipeline can also be used to benchmark new methods in an unbiased manner or to reproduce results obtained from data stored in the Sequence Read Archive (SRA) with an additional dockstore workflow step [37, 38]

To extend the reach of SIMS to investigators without coding experience, we developed a web application based on Streamlit. This application allows users to perform predictions based on pretrained SIMS models. To access the web application the user has to enter the webpage at https://sc-sim s-app.streamlit.app/. Once there, the user has to upload their dataset of interest in h5ad format, select one of our pretrained models and perform the predictions. They will be able to download the predictions in csv format and visualize their labeled data as a UMAP.

### 2.2. Benchmarking SIMS against existing cell classifiers of single cell RNA data

We conducted benchmark tests in three distinct datasets to evaluate SIMS’ performance against other methods built on various theoretical approaches. The first dataset we utilized was the PBMC68K, also known as Zheng68K, derived from human peripheral blood mononuclear cells [39]. This dataset is particularly valuable due to its complex nature, featuring unbalanced cell clusters and cells with similar molecular identities, making it a robust choice for benchmarking cell type annotation methods, as it has been extensively employed for this purpose. As a second dataset we included the human heart dataset, also known as Tucker’s dataset, comprising 11 cell types and exhibiting unbalanced cell clusters [40]. This dataset shares similarities with Zheng68K but contains a significantly larger number of cells (287,000 cells compared to 68,000 cells). Additionally, we incorporated the Human cell landscape, also known as Han’s dataset [18] into our analysis, primarily for its substantial size (over 584,000 cells) and the presence of a wide array of different cell types, totaling 102.

In our benchmarking study, we selected a range of tools that represent diverse methodologies and functionalities within the field of single-cell analysis. The scVI and scANVI pipeline was included owing to their deep learning foundation, utilizing a variational autoencoder to create cell embeddings [27]. This latent representation serves as the basis for subsequent model building and label transfer, making scVI and scANVI essential benchmark for evaluating deep learning-based approaches in single-cell analysis illustrating the scArches package [24]. Another deep learning-based tool, ScNym, adopts an-other two-step process. Beginning with adversarial pretraining, the network is refined through fine-tuning for classification, offering a unique perspective on how deep learning models can be optimized for single-cell RNA data anal-ysis [28]. In contrast, SciBet adopts a non-deep learning approach by fitting multinomial models to the mean expression of marker genes. SciBet was benchmarked primarily for its inference speed, a crucial aspect considering its real-time web-enabled inference capabilities[41]. Seurat, a well-established framework in the field, was included due to its versatility in preprocessing, visualization, and analysis of single-cell data. Additionally, Seurat provides label transfer functionality through the identification of anchors, establish-ing pairwise correspondences between cells in different datasets[19]. We also wanted to evaluate a model with a simpler paradigm behind it, SingleR, which employs a correlation-based method, focusing on variable genes in the reference dataset for calculating differences between cell types. Additionally, an attempt was made to benchmark against scBERT, a large transformer-based model[25]. However, due to its computational complexity, we faced limitations. Despite experimenting with an A10 GPU, scBERT’s demands were such that we were unable to train or evaluate it on any dataset, even with a minimal batch size of 1. These carefully chosen tools enabled a com-prehensive evaluation, considering various approaches and methodologies in the realm of single-cell analysis.

To ensure the robustness of our findings and mitigate the influence of ran-domness, we employed a fivefold cross-validation strategy. Notably, SIMS consistently outperformed the majority of label transfer methods in terms of accuracy (Figure 1; Supplemental Table 1) and Macro F1 score (Supple-mental Figure 5; Supplemental Table 2) across these diverse datasets. This compelling evidence underscores SIMS as a highly accurate and robust clas-sifier, demonstrating its proficiency across diverse tissue types. Additionally, SIMS exhibits scalability to accommodate a large number of cells and show-cases its ability to effectively classify datasets with imbalanced cell types.

**Figure 1:**
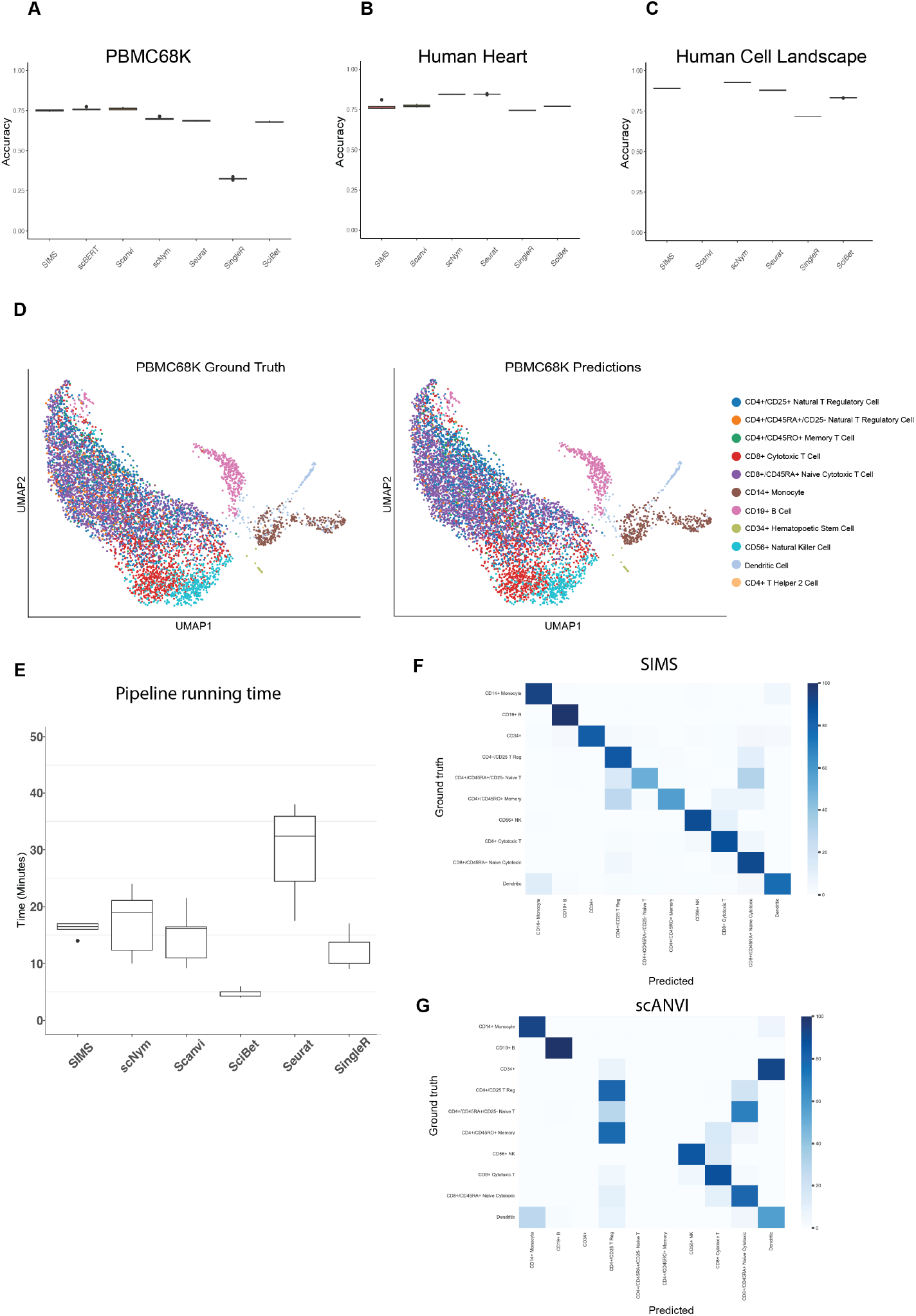
Benchmarking SIMS against other cell classifiers. A)Performance of cell type annotation methods measured by accuracy in the PBMC68K dataset using fivefold cross-validation. Box plots show the median (centre lines), interquartile range (hinges) and 1.5-times the interquartile range (whiskers). B) Performance of cell type annotation methods measured by accuracy in the Human heart dataset. C) Performance of cell type annotation methods measured by accuracy in the Human cell landscape dataset. D) UMAP representation of the PBMC68K cells, colored by ground truth cell type and representation of the PBMC68K cells, colored by SIMS predicted cell type. E) Performance of cell type annotation methods measured by pipeline running time in minutes.F) Heatmap for PBMC68K comparing ground truth annotations and predictions by SIMS G) Heatmap for PBMC68K comparing ground truth annotations and predictions by SCANVI

We also conducted a consistent evaluation of pipeline running times by employing fivefold cross-validation to assess the speed of the benchmarked tools in minutes, using the same comparison methodology (Figure 1E). This analysis was carried out within the NRP clusters[42], leveraging user-accessible GPUs. Whenever feasible, training and inference processes were executed on GPUs; otherwise, they were performed on CPUs.

### 2.3. SIMS accurately performs label transfer in highly complex single cell data: Mouse adult cerebral cortex and hippocampus

Given that SIMS outperforms most state-of-the-art label transfer meth-ods in different datasets, we then asked whether it could perform accurately in a highly complex tissue, such as the brain. We focused in adult mouse cortical and hippocampal data generated by the Allen Brain Institute [43, 44, 45].

The cerebral cortex is among the most complex tissues due to its cellu-lar diversity, the variety and scope of its functions and its transcriptional regulation [46]. The cerebral cortex is organized in 6 layers, and several cortical areas, each with different composition and proportions of excitatory projection neurons (PNs), inhibitory interneurons (INs), glial cells and other non-neuronal cell types [46]. The hippocampus, on the other hand, is part of the archicortex (also known as allocortex) [47]. It is further subdivided into cornu ammonis (CA), dentate gyrus, subiculum, and entorhinal area [47]. While the hippocampus also has a layered structure, made of 3 layers, the cell type composition and numbers vary greatly from those in cerebral cortex [47]. The great diversity of cell types, the close relationship between some of those subtypes, and the anatomical separation between these regions, make cere-bral cortex and hippocampal datasets complex but attractive benchmarking models to test SIMS.

The dataset contained 42 cell types, including PNs, INs, endothelial and glia cells. Training in 80% of the cells selected at random and testing on the remaining 20%, we find that SIMS performs at an accuracy of 97.6% and a Macro F1 score of 0.983 (Figure 2 and Supplemental Figure 6).

**Figure 2:**
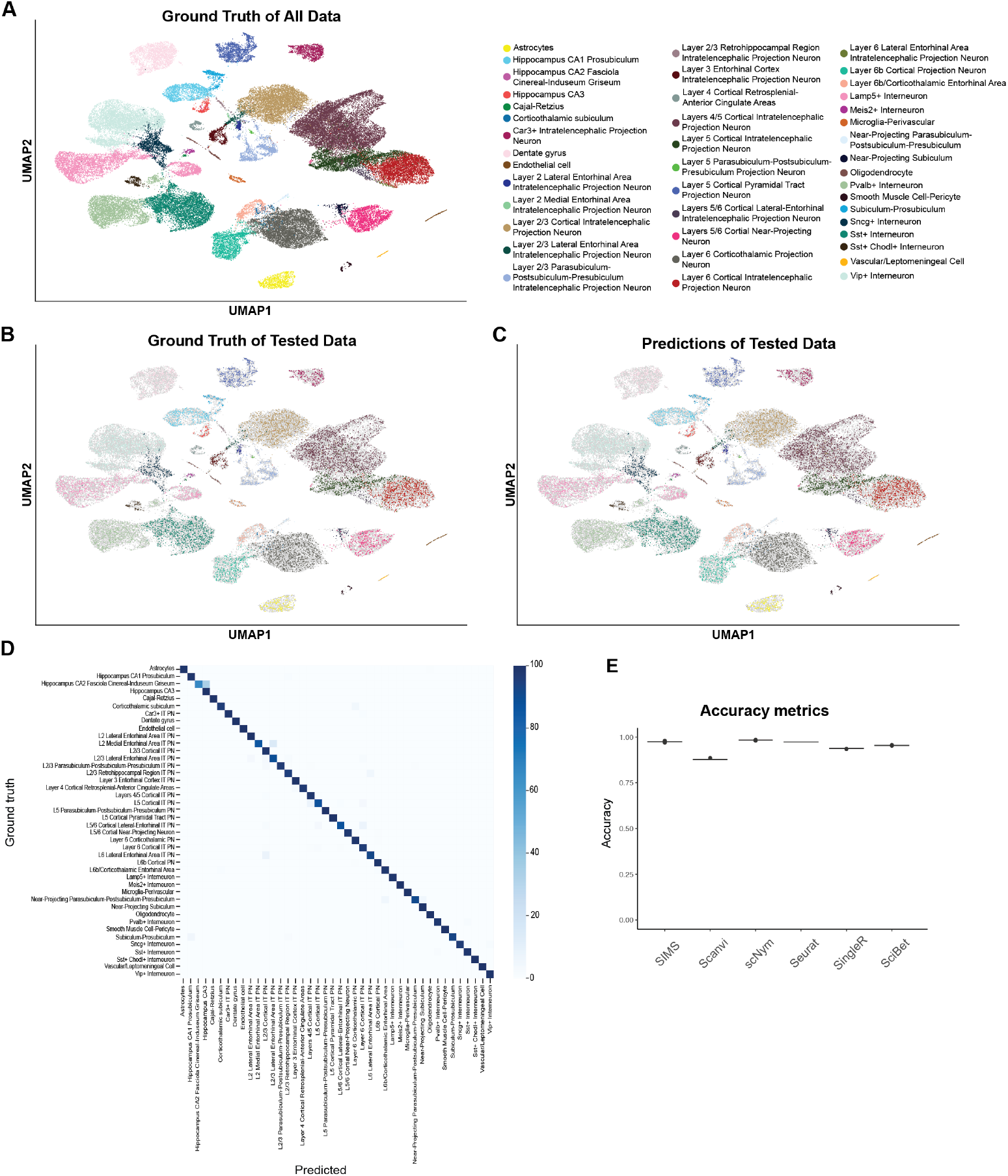
Application of SIMS to single Cell RNA sequencing: Adult Mouse Cerebral Cortex and Hippocampus. A) Ground truth UMAP representation for the dataset. B) Ground truth UMAP representation for the Subset of Cells used for testing the algorithm in the train-test split. C) Predictions made by SIMS in that subset of data. D) Confusion Matrix for the test-split. L= Layer; IT = Intratelencephalic; PN = Projection Neuron. E) Benchmarking SIMS against other cell classifiers. F) Performance of cell type annotation methods measured by accuracy in the Allen mouse dataset using fivefold cross-validation. Box plots show the median (centre lines), interquartile range (hinges) and 1.5-times the interquartile range (whiskers)

We then performed ablation studies to investigate the performance of SIMS. We find that training in as little as 7% of the dataset (3,285 cells) is sufficient to obtain a label transfer accuracy of over 95% and Median F1 score of over 0.95 (Supplemental Figure 7). The Macro F1 after training in 7% of the data is 0.90 (Supplemental Figure 7). Given the low amount of training data needed to obtain a high accuracy in label transfer, we conclude that SIMS is a data efficient machine learning model.

SIMS provides interpretability by computing weights for sparse feature masks in the encoding layer. These weights indicate the most influential genes in the network’s decision-making for assigning cell types. To assess this interpretability, we generated three dataset partitions with varying levels of granularity. Our aim was to observe if the network could accurately select pertinent genes to distinguish the groups formed at each resolution level. In order to analyze the results we focused in the Pvalb+ INs, a group of inhibitory neurons born in the Medial ganglionic eminence (MGE). For the lowest level of granularity, which limit the cell options to INs, PNs and Non-Neuronal Cells, we find that for the INs group some important genes selected by the model were Kcnip and Igf1 (Figure 3A-B), both of which have been previously shown to be important IN genes [48, 49, 50]. For the medium level of granularity (Medial ganglionic eminence, non medial ganglionic eminence), and consistent with previous literature we find that for the MGE-derived INs the genes selected were Rpp25, Dlx1, Dlx5, Gad1, Ffg13 and Cck. [51, 50, 52, 53] (Supplemental Figure 8). For the highest level of granularity (Pvalb+ INs), some of the selected genes were Satb1, Pvalb, Lypd6, Dlx6os-1 and Bmp3. [53] (Figure 3C-D)

**Figure 3:**
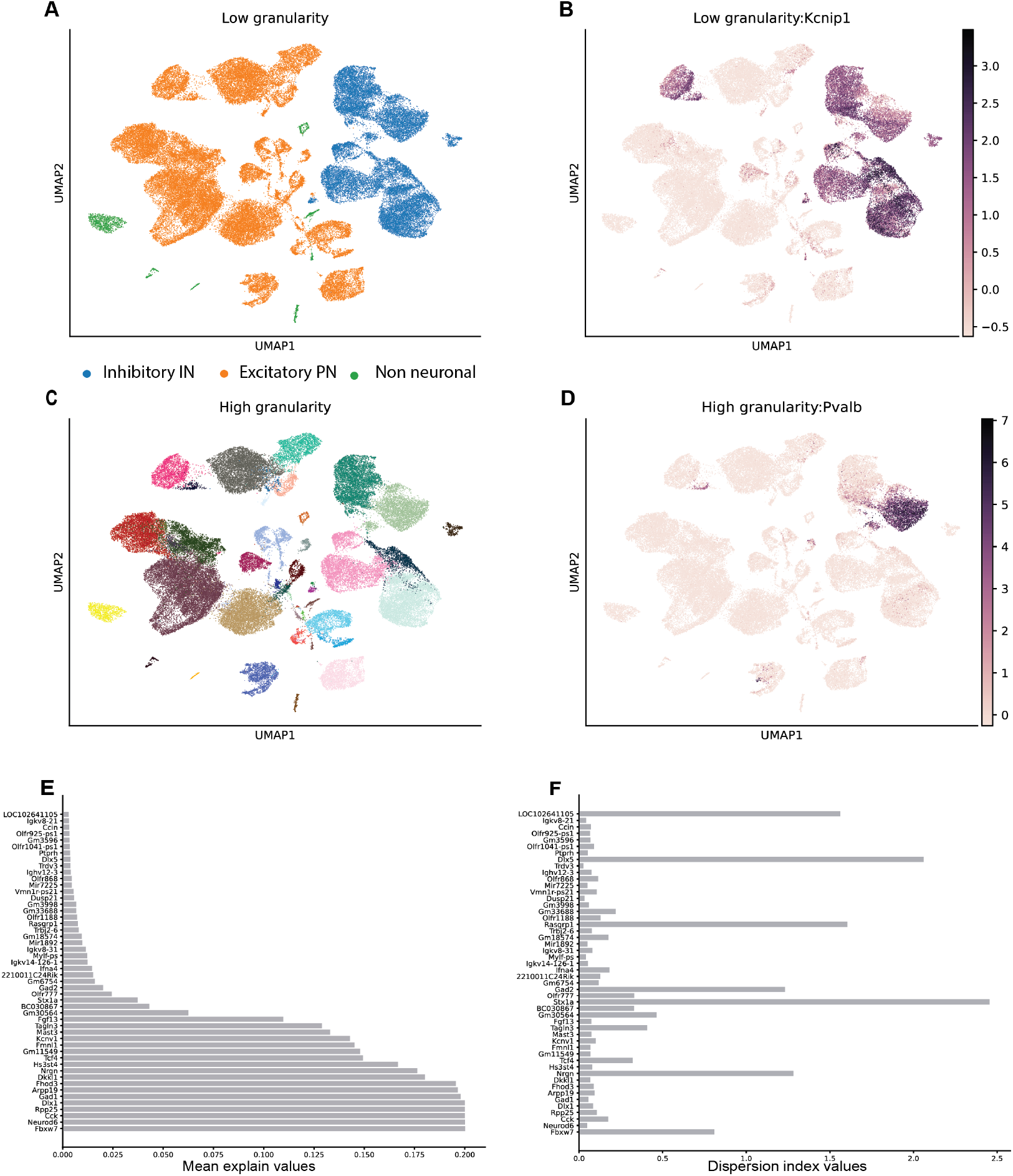
SIMS explainability. A) UMAP representation of the Allen Mouse dataset coloured by macro cell type . B) UMAP representation of the Allen Mouse dataset coloured by expression of the selected gene by SIMS for the GABAergic group. C) UMAP representation of the Allen Mouse dataset coloured by cell type. Same naming convention used for figure 2A. D) UMAP representation of the Allen Mouse dataset coloured by expression of the selected gene by SIMS for the PVALB+ interneuron group. E) Mean explain value for the top 50 genes across 300 runs. F) Dispersion index value for the top 50 genes across 300 runs.

To confirm that the selection of the most important genes was consistent across different runs we performed the experiment with the highest level of granularity 300 times. For each experiment we normalized each gene weight against the highest weight gene measured in that run and measured the mean weight and dispersion index for each gene across all runs (Figure 3E-F). Given the explainability matrix *E ∈* R*^n×m^* comprised of *m* genes measured across *n* cells, we select all rows representing cells with the same predicted label and compute:

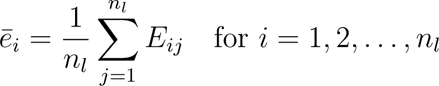

We then average *ē_i_* across all 300 runs. To calculate the dispersion index, we first measured the average importance of each gene across all 300 runs

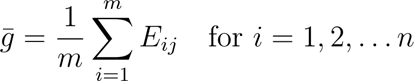

and then compute the dispersion index as

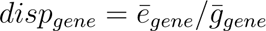

In the top 10 of genes more important for classification we can find Excitatory PN markers (Neurod6), Inhibitory IN markers (Cck, Rpp25, Dlx1, Gad1), neural progenitor related genes (Fbxw7) and genes related to different neuropsychiatric disorders (Arpp19, Fhod3, Nrgn). Top genes show mean explain values around 0.2 (Figure 3E), for comparison the mean explain value for the median gene is around 10*^−^*^6^ (Supplemental Figure 10). This showcases the consistency of gene selection by SIMS and how it could be used to find clinically relevant genes overlooked by conventional methods.

### 2.4. SIMS accurately performs trans-sample label transfer in highly complex single nuclei data: Human adult cerebral cortex

Single nuclei RNA sequencing has become an important emerging tool in the generation of atlases, particularly in tissues where obtaining single cells is difficult. Cell nuclei are used in neuroscience because adult neurons are difficult to obtain, due to their high connectivity, sensitivity to dissociation enzymes and high fragility, often resulting in datasets with abundant cell death, low neuronal representation and low quality RNA [54]. Importantly, single nuclei sequencing is compatible with cryopreserved banked tissue [55]. Yet, the data generated in single nuclei RNA sequencing is not necessarily similar to the data generated in single cell RNA sequencing. For instance, a recent study comparing the abundance of cell activation-related genes in microglia sequenced using single cell and single nuclei technologies, showed significant differences between both datasets [56]. Moreover, single nuclei datasets are more prone to ambient RNA contamination from the lysed cells [57]. In the case of the brain, it has been observed that neuronal ambi-ent RNA has masked the transcriptomic signature of glia cells, leading to incorrect classification of glia subclasses in existing atlases [57].

Given the high label transfer accuracy of SIMS in single-cell data, we then tested its performance in single nuclei datasets. As a proof of principle, we selected the human adult cerebral cortex dataset generated by the Allen Brain Institute [44, 43]. We trained on 80% of the data and tested the model in the remaining 20%. Overall, we obtained an accuracy: 98.0% and a Macro F1-score of 0.974 (Figure 4; Supplemental Figure 9; Table 1).

**Figure 4:**
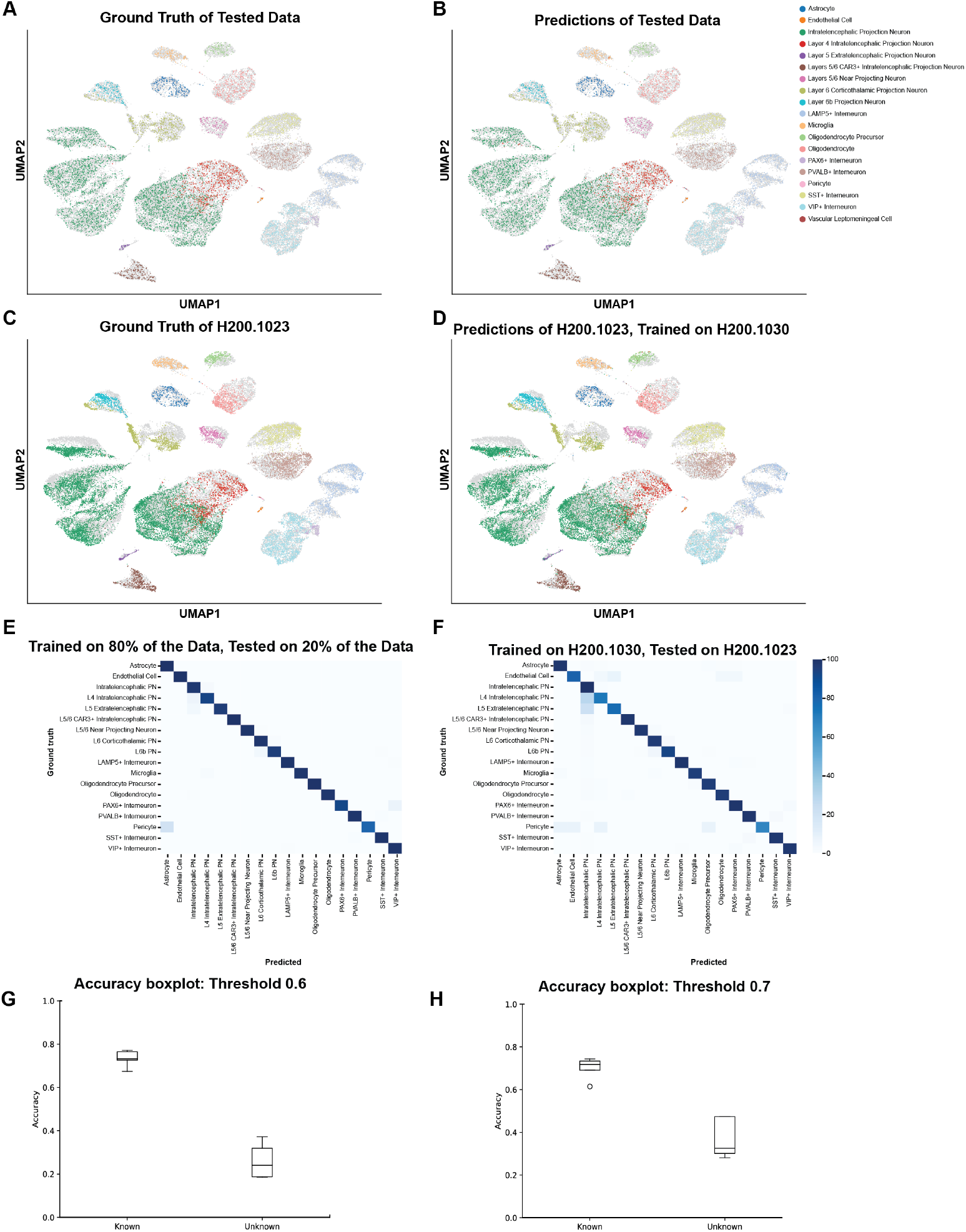
Application of SIMS to trans-sample predictions of single Nuclei RNA sequencing: Adult human cerebral cortex. A) Ground truth for the test-split data. B) Predictions for the test-split data. C) Ground truth for the H200.1023 sample. D) Prediction for the H200.1023 sample after training on the H200.1030 sample. E) Confusion matrix for the test Split. F) Confusion matrix for the test Split. G) Accuracy boxplot for the Known and Unknown cell classification with a confidence threshold of 0.6 H) Accuracy boxplot for the Known and Unknown cell classification with a confidence threshold of 0.7. L = Cortical Layer; PN = Projection Neuron. Additional examples are on Supplemental Figure 12.

**Figure 5:**
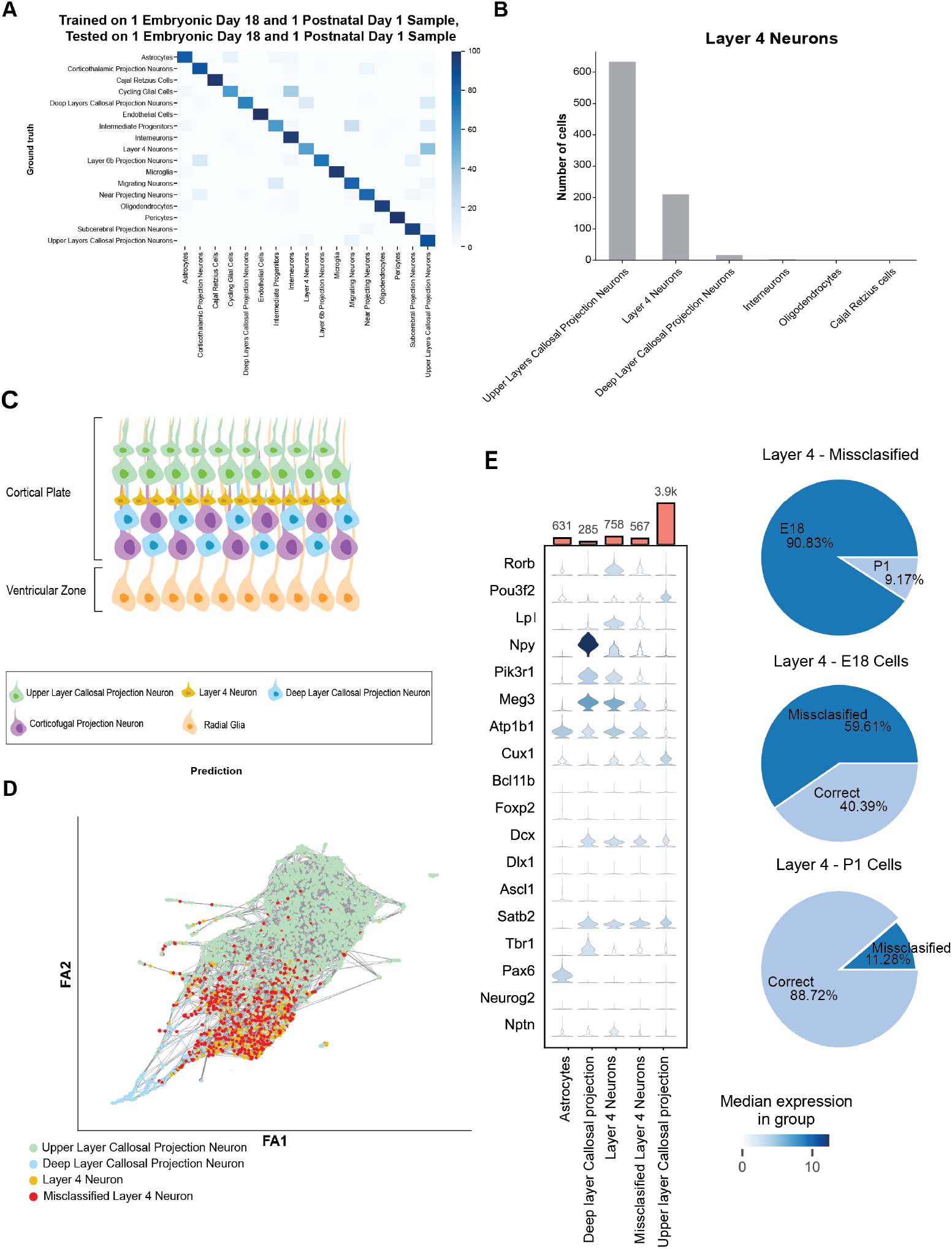
Application of SIMS to developing tissue: Mouse cerebral cortex. A) Confusion Matrix for E18P1 split, where we trained on Sample 1 E18 and Sample 1 P1 and predicted on Sample 2 E18 and Sample 2 P1 B) Barplot showing the number of Layer 4 Cells that get predicted as the C) Diagram of the mouse cerebral cortex after neurogenesis. D) Force Atlas representation of Layer 4 Neurons. E) Violin plot showing gene expression in the misclassified Layer 4 group compared to the groups that is classified as.

**Table 1:** Trans-sample accuracies and Macro F1-scores for human adult cere-bral cortex dataset.

| Training Sample | Testing Data | Accuracy | Macro F1-score |
| --- | --- | --- | --- |
| 80% of Data | 20% of Data | 98.0% | 0.974 |
| H200.1023 | H200.1025 | 94.0% | 0.84 |
| H200.1023 | H200.1030 | 94.4% | 0.865 |
| H200.1025 | H200.1023 | 93.1% | 0.769 |
| H200.1025 | H200.1030 | 93.1% | 0.779 |
| H200.1030 | H200.1023 | 95.8% | 0.862 |
| H200.1030 | H200.1025 | 94.8% | 0.87 |

We then performed a data ablation study and observed that we obtained over 95% accuracy using as little as 7% of data for training (2,124 cells). Similarly, we obtained a Macro F1-score of over 0.95 with 9% (2,731 cells) of the data and a median F1 of over 0.95 with 8% of the data (2,428 cells) for training (Supplemental Figure 11).

We then asked how SIMS performs in trans-sample predictions. This dataset is made of 3 different postmortem samples. Namely: H200.1023, a 43 years old Iranian-descent woman; H200.1025, a 50 years old Caucasian male; and H200.1030, a 57 years old Caucasian male. We trained the model on one sample and tested it on the other 2 samples. We performed this experiment in each possible combination, obtaining accuracies ranging from 93.1 to 95.8% (Figure 4; Supplemental Figure 12; Table 1; Supplemental Tables 3-8).

As shown, SIMS predicts the label accurately for most cell types across samples. SIMS shows a decrease in performance when trying to classify Pericytes as sometimes it labels them as Astrocytes (Supplemental Tables 3-8). This is consistent with recent work showing that previously annotated single nuclei atlases of the brain often mask non-neuronal cell types [57]. In addition, we observed that Layer 4 Intratelencephalic neurons often get classified as generic Intratelencephalic neurons (Supplemental Tables 3-8). This is in agreement with the fact that Layer 4 Intratelencephalic neurons are a subset of Intratelencephalic neurons [58]. We also employed this dataset to assess the capacity of SIMS to differentiate between recognized cell types and those not included in the training dataset. This capability holds significance as it can function as a surrogate metric for identifying cells in new datasets that were absent from the reference dataset used for training. In this particular scenario, we implemented a leave-one-out methodology, where we excluded one cell type from the training dataset and then made predictions on the test set, encompassing all of its cell types. Subsequent to temperature scaling, we utilized the model’s probability outputs as a measure of confidence, such that a probability of 0.5 approximately measures that the model possesses a 50% level of confidence in the predicted cell type’s accuracy. Following this, we established a user-adjustable threshold to determine whether the cell type should be labeled as the predicted cell type or categorized as an unknown cell type (Figure 4G-H). Altogether, we conclude that SIMS is a powerful approach to perform intra-sample and trans-sample label transfer in complex and highly diverse tissues such as the adult brain.

### 2.5. SIMS can accurately classify cells during neuronal specification

Having established that SIMS can accurately predict cell labels in complex tissues, we then asked how our model performed predicting cells of different ages. Classifying cells during development is challenging, as several spatiotemporal dynamics can mask the biological cell identities [59]. During cortical development, gene networks of competing neuronal identities first colocalize within the same cells and are further segregate postmitotically [60, 46, 61], likely through activity-dependent mechanisms [62, 63].

To test the accuracy of SIMS at classifying developing tissue, we focused on mouse cortical development due to its short timeline [64]. In the mouse cortex, neurogenesis starts at embryonic day (E) 11.5, and it is mostly completed by E15.5 [64]. Common C57BL/6 laboratory mice are born at E18.5 [65]. Neonatal mice are timed based on the postnatal day (P) [65]. We took advantage of a cell atlas of mouse cortical development that contains 2 samples of E18 mouse embryos and 2 samples of P1 mice [60]. These timed samples, which are 1 day apart from each other represent timepoints at which all mouse neurogenesis is completed [64]. At these timepoints, neurons may still be undergoing fate refinement [66], and consequently retain fate plasticity, albeit limited [67, 68, 69].

First, we trained a model on one E18 and one P1 sample and tested the accuracy of label transfer in two samples, one of each age (Supplemental Figure 13 A-B). Across 17 cell types, we find that the model predicts the labels with an accuracy of 84.2% and a Macro F1-score of 0.791 (Figure 5A; Supplemental Table 9).

We then tested SIMS by training on two P1 samples and testing the label transfer in two E18 samples (Supplemental Figure 13 C-D). We find that in this experiment, the label transfer accuracy drops to 73.6% and the Macro F1-score to 0.674 (Figure 5B; Supplemental Table 10). Interestingly, however, this drop in accuracy is not random, but either follows the developmental trajectories of the misclassified cells or misclassifies cells as transcriptomically similar cell types. For example, astrocytes are a subtype of glia cells that retain the ability to divide throughout life [70]. Indeed the major source of astrocytes in the cerebral cortex is other dividing astrocytes [70]. Consequently, the “Cycling Glia Cells” cluster is often predicted as astrocytes (Supplemental Figure 13). In the neuronal lineage, we find that SIMS can accurately predict most cell types. Going back to the combined ages model, we focused on Layer 4 neurons, which is one of the neuronal subtypes with the lowest accuracy in label transfer (24.31%). We find that these neurons are often classified as upper layer callosal PNs, and rarely as callosal PNs of the deep layers (Figure 5B-E). While morphologically distinct, layer 4 neurons share transcriptional homology with callosal PNs [60, 71]. Indeed, recent work has shown that Layer 4 neurons transiently have a callosal-projecting axon, which is postmitotically eliminated during circuit maturation, well after P1 [58]. In agreement, Layer 4 neurons that are mislocalized to the upper cortical layers retain an upper layer callosal PN identity and fail to refine their identity [72]. By comparing the gene expression of upper layer callosal PNs, the correctly classified Layer 4 neurons and the misclassified Layer 4 neurons, we observe that while upper layer callosal PNs and correctly classified Layer 4 neurons have the gene expression patterns proper to their identity, misclassified Layer 4 neurons have an intermediate expression of genes that define the identity of the other two cell types, such as Rorb[73] (Figure 5). Notably, most (90.1%) of the misclassified Layer 4 neurons belong to the E18, likely representing neurons undergoing fate refinement. Altogether, this example highlights the difficulty that cell classifiers face when trying to discretely label cells during development.

Together, we conclude that SIMS can accurately predict cell labels of specified neurons. However, when applying SIMS during periods of differentiation and fate refinement, it uncovers similar identities in the developmental trajectories. This is likely caused by transcriptomic similarities that can often mask the proper identification. Alternatively, SIMS may identify subtle differences in fate transitions that cannot be accurately pinpointed by traditional clustering methods in the reference atlases.

### 2.6. SIMS identifies cell-line differences in gene expression in human cortical organoids

Cortical organoids are a powerful tool to study brain development, evolution and disease [13, 74, 75]. Yet, like many pluripotent stem cell-derived models, cortical organoids are affected by cell line variability and culture conditions that can affect the reproducibility of the protocols [76]. Moreover, transcriptomic analysis of cortical organoids has revealed strong signatures of cell stress [77, 78, 79], which can impair proper cell type specification [80]. In addition, *in vitro* conditions generate cell types of uncharacterized identity, that do not have an *in vivo* counterpart [78, 81]. While some have argued that these cells should be removed from further analysis [81], the most common approach is to annotate them as “Unknown” cell clusters [74].

To understand whether SIMS could be used to uncover cell line differ-ences and identify different trajectories, we used a dataset from 6 months old human cortical organoids derived from 3 different cell lines (3 organoids per batch), each with their own idiosyncrasy [74]. Specifically, this dataset contained: 1) one batch of cortical organoids derived from the 11A cell line, in which all cells had been identified and no cell was labeled as “Unknown”, 2) one batch of cortical organoids derived from the GM8330 cell line, which contained a small number of “Unknown” cells and a large proportion of Im-mature INs, and 3) two batches of cortical organoids derived from the PGP1 cell line, which contained major batch effects. One of those batches had a large number of “Unknown” cells and cells of poor quality, and was therefore dropped from further analysis (Figure 6A-B; Supplemental Figure 14).

**Figure 6:**
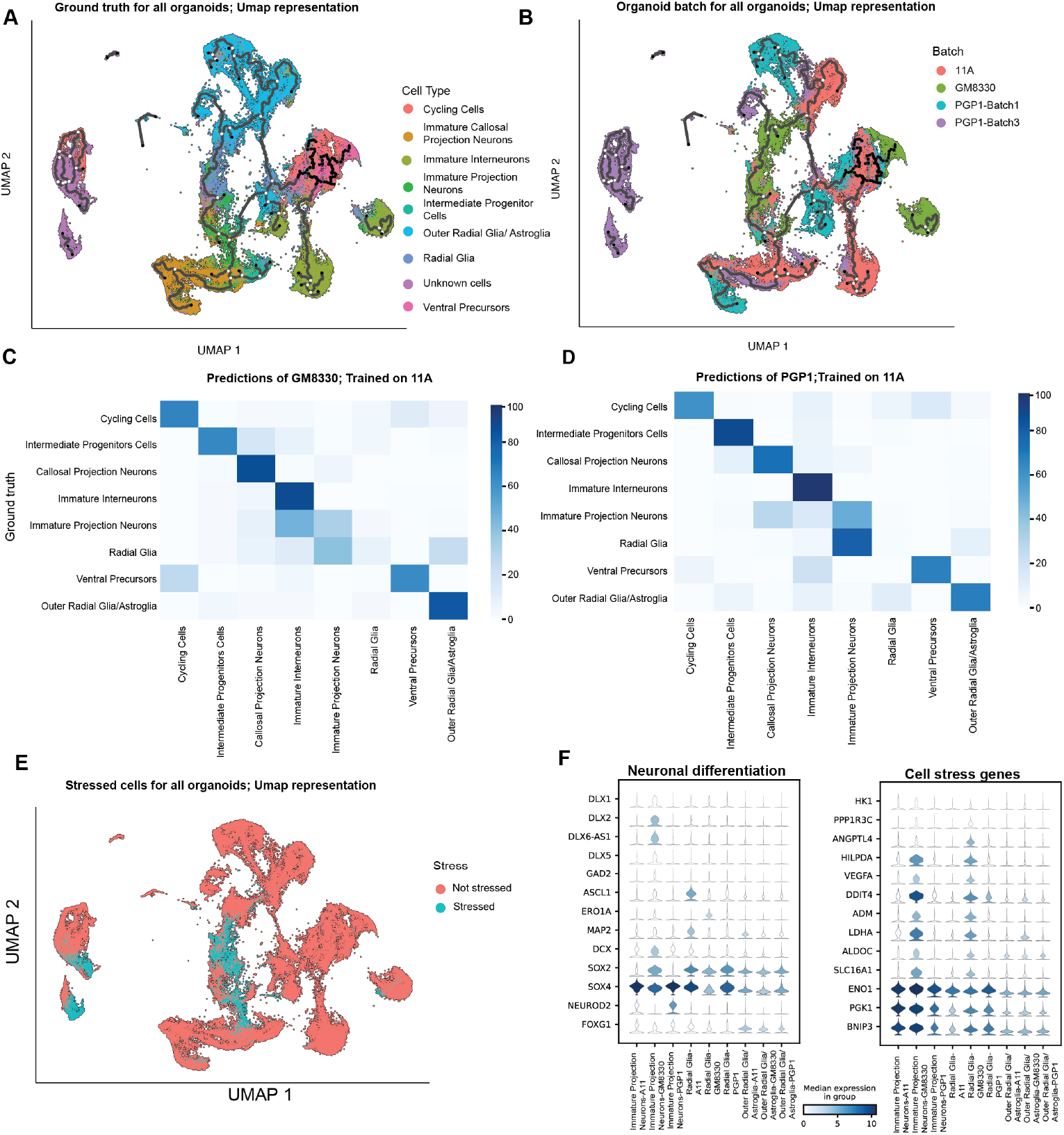
Application of SIMS to *in vitro* generated models: hu-man cortical organoids. A) UMAP representation of the Ground truth cell type for all cell lines. B) UMAP representation of the batch and cell line for all cell lines C) Confusion Matrix for GM3880-derived organoids, model trained on 11A-derived organoids. D) Confusion Matrix for PGP1-derived organoids, model trained on 11A-derived organoids.E) UMAP representation for stressed cells as annotated by Gruffi in all organoids. F)Violin plots for neuronal differentiation and Cell stress genes showing differences among cell lines

We performed label transfers between organoids generated from the three cell lines. We first performed an intra-cell line label transfer using the 11A organoids. We trained on 2 organoids and predicted the cells on a third organoid. We find an Accuracy of 86.0% and a Macro F1-score of 0.794 (Sup-plemental Figure 15). We then performed trans-cell line predictions training on 11A and predicting the cell types of the other lines. We obtained an Ac-curacy of 71.3% and a Macro F1-score of 0.564 when predicting cells from PGP1 organoids and an accuracy of 67.4% and a Macro F1-score of 0.570 when predicting cells from GM8330 organoids. We observe a high degree of accuracy for most cell types tested, including Cycling Cells, Intermedi-ate Progenitor Cells, Outer Radial Glia/Astroglia, Immature INs, Ventral Precursors and Callosal PNs (Supplemental Table 11). Interestingly, Radial Glia cells (RGs) from both PGP1 and GM8330 cell lines often were classified as Immature PNs. Specifically, we find that 82% of the PGP1 and 42% of the GM8330 RGs get predicted as Immature PNs when the data is trained on the 11A cell line (Figure 6C-D). Strikingly, only 1.9% of PGP1 RGs and 3.9% of GM8330 RGs are predicted as RGs. These results suggest major differences in gene expression between the RG annotated cells across cortical organoids derived from different cell lines.

Previous work has shown that cell stress in organoids impairs proper fate acquisition of PNs [80]. We therefore took advantage of Gruffi, a recently de-veloped tool to annotate stressed cells in human neuronal tissue [81]. Overall, we find that organoids derived from the GM8330 cell line showed the biggest percentage of stressed cells (16.67%), while organoids derived from the PGP1 and 11A cell lines had 6.6% and 4.9% of stressed cells, respectively.(Figure 6E). To understand whether the stressed cells were responsible for the mis-classfication, we removed these cells from the 11A training set. We then performed a new round of label transfers. Using this approach, we find that 56% of PGP1-derived RGs and 27%-derived RGs continue to be classified as Immature PNs. Importantly, only 7.2% of PGP1-derived and 14% of GM8330-derived RGs are predicted as RGs.

We then removed the stressed cells from both the training and the predicted datasets and find that 44% of PGP1-derived and 14% of GM8330-derived RGs are classified as Immature PNs. Notably, the number of RGs that are properly classified as such remains similar, with only 6.9% of PGP1-derived and 19% of GM8330-derived RGs properly predicted. Altogether, these results suggest that cell stress alone cannot explain the differences in cell expression between RGs of cell lines.

### 2.7. SIMS identifies improperly annotated cell lineages in human cortical organoid atlases

Given that label transfer between human cortical organoids derived from different cell lines poorly predicted the RG cell type, we then focused on assessing the most common predictions for this cell type after stressed cells were removed from both the training and the prediction datasets. While in the PGP1 line the majority of the misclassified RGs are Immature PNs, the second most common cell prediction is the closely related Outer Radial Glia/Astroglia cell type. On the other hand, for the GM8330 cell line the most commonly predicted cell type is Immature INs. Unlike RGs, Outer Radial Glia/Astroglia and Immature PNs that belong to the dorsal telencephalic lineage, INs are derived from the distinct and distant ventral telencephalon [46]. A deeper analysis into the GM8330 cell line reveals that 65% of the Immature PNs also get predicted as Immature INs (Figure 6C), indicating a consistent misclassification between neuronal lineages in the GM8330 cell line. We then performed a Wilcoxon test rank for differential expression analysis between the three cell lines. We found that, unlike the other cell lines, Immature PNs derived from GM8330 organoids expressed genes from the DLX family, present in INs and not in the PN lineage [82] (Supplemental Figure 16). Together, these results suggest an off-target ventralization of organoids derived from the GM8330 cell line.

To confirm this discovery we performed a label transfer experiment training on fetal tissue derived from gestational weeks (GW) 14-25 human embryos [83]. Most cell types, such as cycling cells and ventral precursors get classified as expected. Focusing on neuronal cell types, the majority of Callosal PNs get classified as Excitatory PNs (80% PGP1, 60% GM8330, 74% 11A) and Immature INs are properly classified as INs (93% PGP1, 86% GM8330, 86% 11A). However, Immature PNs have clear difference between the 3 cell lines: For the 11A line, 34% of Immature PNs get classified as Excitatory PNss and 38% as RGs. Similarly, in the PGP1 line, 57% of Immature PNs are classified as Excitatory Ps and 20% as RGs. On the other hand, only 7% of the GM8330 Immature PNs are classified as Excitatory PNs, and 21% are classified as RGs. Importantly 44% of these cells are predicted as INs. (Sup-plemental Figure 17), further suggesting a ventralization of the organoids derived from the GM8330 line.

We then performed a pseudotime analysis using Monocle 3 [84]. In the 11A and PGP1 lines, we observe a clear differentiation trajectory from RG to the Excitatory PN lineage(Immature PNs and Callosal PNs). In these lines, the IN lineage follows a separate path (Figure 7A; Supplemental Figure 18). Focusing on the GM8330 cell line, we observe that a large subset of Immature PNs unexpectedly appear together with the IN lineage (Supplemental Figure 18). Altogether, the data suggests that SIMS has correctly identified that a large subset of cells labeled as Immature PNs in the GM8330 are in fact INs.

**Figure 7:**
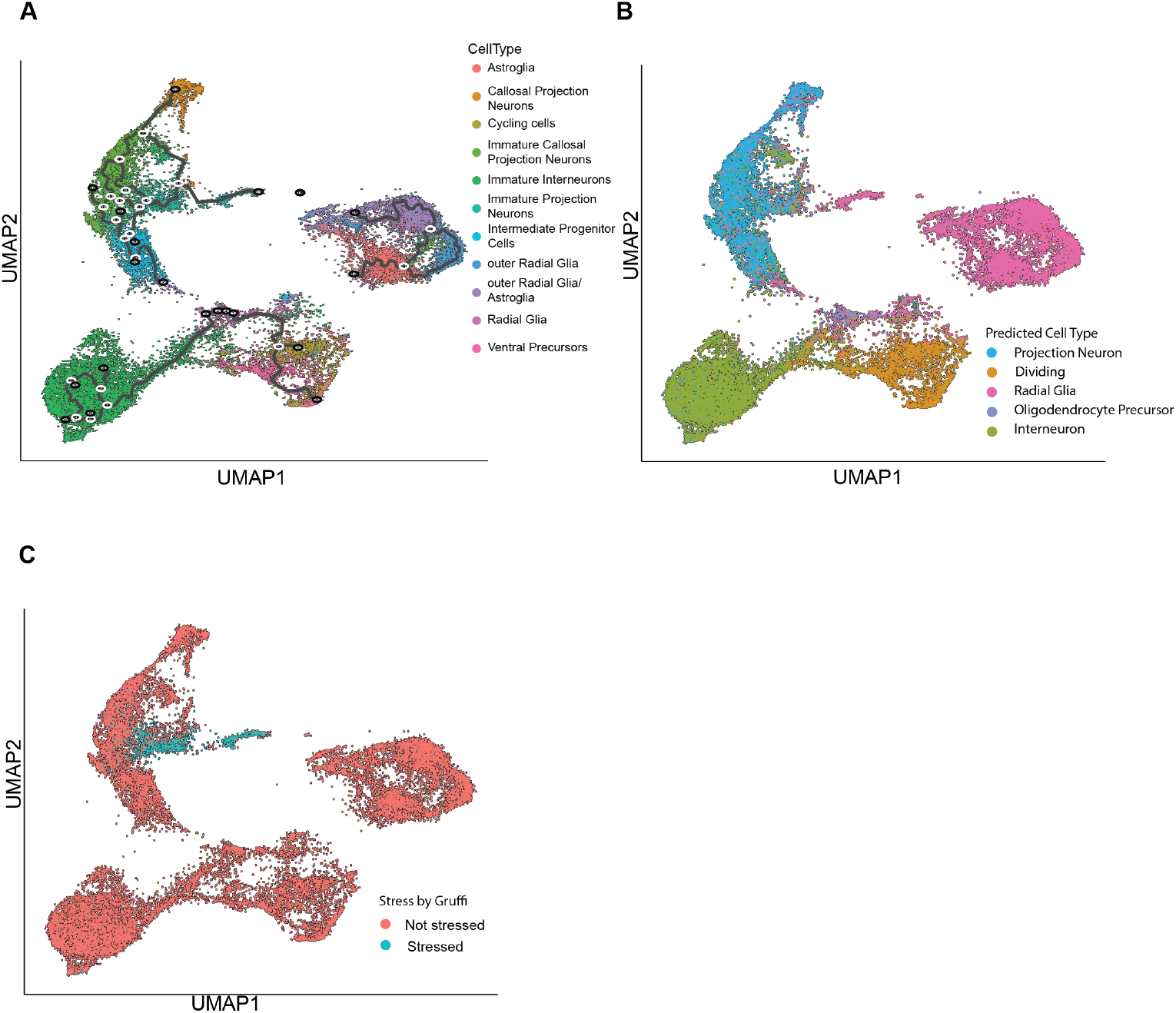
Application of SIMS to *in vitro* generated models: human cortical organoids. A) UMAP representation of the Ground truth cell type for 11A organoids. B) UMAP representation of the label transfer from Fetal tissue for 11A organoids. C) UMAP representation for stressed cells as annotated by Gruffi in the 11A organoids.

### 2.8. Leveraging In Vivo Data Refines Cell Type Prediction in Brain Organoids

Visualization methods based on dimensionality reduction, such as prin-cipal component analysis (PCA) and t-distributed stochastic neighbor em-bedding (tSNE) often miss the global structure of the data and can lead to misclassification of cells [85]. Given that SIMS identified a ventralization of the GM8330 cell line (Figure 6), we then asked whether it can identify other cells previously misclassified in existing atlases [74]. We analyzed 6 months old organoids derived from the 11A cell line. We first performed pseudotime analysis and found that a subset of cells labeled as Immature PNs cluster in between other Immature PNs and Glia Cells (Figure 7A). Interestingly, all these cells are identified by Gruffi as stressed cells (Figure 7B). To test whether these cells were mistakenly classified in previous atlases, we performed a label transfer from GW14-25 primary fetal tissue [83]. We find that SIMS assigns the entirety of this cell cluster as RGs and not PNs (Figure 7C). Gene expression analysis of molecular markers of RGs, such as SOX2 and PAX6 (Supplemental Figure 19), confirm that the SIMS label is correct. In complement, these cells lack expression of PN subtypes markers such TBR1, SATB2, CUX1, CUX2, as well as Pan-PN markers EMX1, DCX, NEUROD2 and NEUROD6 (Supplemental Figure 19). Altogether, these re-sults suggest that the stressed cells previously labeled as Immature PNs in the 11A cell line are indeed RGs.

We asked how correcting the cell type annotation in the 11A affected the label transfer between organoids derived from different cell lines. We trained SIMS in the newly annotated 11A dataset and made predictions in both the PGP1 and the GM8330 cells. We found that for the new model trained on the 11A cell line there is an Accuracy of 75.7% and a Macro F1-score of 0.583 for PGP1 organoids and an Accuracy of 76.3% and a Macro F1-score of 0.603 for GM8330 organoids (Supplemental Table 13,14), representing a significant improvement from label transfer experiments before the reclassification (Sup-plemental Table 11,12). Furthermore, we find that RGs now get predicted at an Accuracy of 43.0% for PGP1 and 32.0% GM8330, as compared to the original predictions of 1.9% and 3.9% for the respective cell lines. Together, we show that proper identification of cell types through label transfer from primary tissue can help systematize multi-sample cell atlases.

## 3. Discussion

Currently, over 1.5M cells per month are sequenced and archived through the different cell atlas projects [86]. With the lowering trends in sequencing costs the number of cells sequenced is increasing exponentially [3, 86]. Yet, cell annotation remains a highly manual process, which is limiting the repro-ducibility and introducing biases in the data. Several open access solutions have emerged to streamline the process, albeit with different accuracies [2].

Deep learning approaches that apply transformer-based architectures to gene expression data have been shown to outperform other commonly used methods [25]. However, these approaches require large number of cells for pretraining their algorithms and advanced computational knowledge and re-sources to further train their models [25]. SIMS does not require pretraining, therefore avoiding large data files and increasing its versatility. An added advantage to SIMS is the requirements with which the training can be per-formed, which allows for the users to run the program in their local comput-ers.

We designed SIMS as a low code tool for both training and perform-ing label transfer across single cell datasets (Figure 1). SIMS can be used on user-specified datasets, rather than reference datasets that are usually a prerequisite in popular tools. This is meant to remove barriers in adoption by new labs, medical practitioners, students and non-experts alike. Unlike other deep learning models [25], SIMS can use genes that are defined by the user, allowing the label transfer in novel genomes, or use annotated genomes without standard nomenclature. Other deep learning approaches, such as scBERT [25], have been shown to work well with datasets of up to 16K genes. SIMS, being based on TabNet, and therefore optimized for tabular data [30], can work well with over 45K features (Figure 2). This property would allow, in principle, SIMS to be trained simultaneously on references of multiple species, species with large genomes such as the axolotl [87], as well as multimodal data including combined single cell gene expression and gene accessibility sequencing datasets [88].

When it comes to interpretability SIMS is able to output a sparse selection of the most important genes, that can then be easily plotted in the Python ecosystem of Scanpy, while other tools [25] rely on external cross-platform packages. This can hamper the adoption of new users, including non bioinformaticians [89]. Indeed, non-experts could greatly benefit from intuitive and low effort tools that can streamline the analysis and integration of their newly generated data with existing knowledge [89]. To facilitate its adoption, we created a web app and a Terra pipeline that can be easily adopted with minimal coding knowledge and low infrastructural resources, offering accesible cloud computing. Furthermore, our approaches facilitate the sharing of trained models which can streamline collaboration between multiple groups.

After showing that SIMS performs as good or better than state of the art methods, we focused on applying this tool to data generated from the brain. The brain is a complex tissue, where the great diversity of neurons is generated over a relatively short time period and identities are refined throughout life [46, 66]. Several efforts, such as the BRAIN Initiative and others, exist to sequence neurons across different ages, species, and diseases [90, 91]. While the neuroscience community has started efforts to agree on naming conventions across the increasing number of datasets [5, 92], there is still significant ontological inconsistencies in existing publications. We believe that SIMS could become an important tool to streamline these community-driven efforts. It is important to mention that while we focused our work in the brain, SIMS can easily be applied to single cell RNA sequencing data of any other organ.

When performing label transfer in fully differentiated neuronal cell types, SIMS performed remarkably well, with accuracies above 97%. Unlike many other tools, which define cells by the strong expression of marker genes [7, 93], the SIMS model takes advantage of lack of expression, and fluctuations of expression levels of the whole transcriptome to learn and identify cell labels. Consistent with this, we observed that in developing tissue, where gene expression is fluctuating and identities are being refined, SIMS was able to classify most cell types and identify maturation differences in cell types undergoing fate refinement.

When applied to cortical organoids, SIMS identified previously misan-notated cells in existing atlases [74].These errors in annotation were caused by traditional clustering followed by differential gene expression analysis and marker identification [74]. Notably, stressed cells were often misannotated, which is a common issue in organoid development [80, 81]. Revisiting and re-annotating existing atlases will greatly increase the accuracy of label transfer and improve the development of future protocols. Furthermore, annotating stem cell-derived atlases using primary fetal samples as reference can be used as a gold standard in the field and to discover cell types underrepresented in the existing protocols [74, 91].

Applying SIMS to developing brain tissue including primary samples and organoids, allowed us to identify subtle differences in developmental trajec-tories between cell types generated. We therefore believe that SIMS can be of great value at studying developmental disorders, such as Autism, where existing models have already shown cell-type dependent asynchronous de-velopmental trajectories in different neuronal lineages [94]. Hybrid pipelines that integrate pseudotime-focused tools, such as Monocle or BOMA [84, 7], could become complementary to SIMS and have the potential to provide more comprehensive insights into these questions.

While we have shown that SIMS can accurately predict trans-sample labels and perform label transfer across different methodologies (single cell and single nuclei RNA sequencing) and models (primary tissue and cortical organoids), we have limited our work to samples within the same species. This is because neuronal subtypes diverge significantly between species [44] and at the individual level gene orthologs can show different expression levels in different species [95]. However, some neuronal subtypes, such as MGE-derived INs, are transcriptomically more conserved across evolution than other primary neurons, including cortical PNs [13, 44]. In the future, these IN subtypes could be used as a way to validate SIMS to perform trans-species predictions [96]. Additional modifications, such as gene module extraction could provide increased accuracy for label transfer, as meta-modules could prove to be more conserved between evolutionary distant species than gene orthologs [92, 97, 98].

In conclusion, we propose SIMS as a novel, accurate and easy to use tool to facilitate label transfer in single cell data with a direct application in the neuroscience community.

## 4. Material and methods

### 4.1. The SIMS Pipeline

The classifier component of the SIMS framework is TabNet [30], a transformer-based neural network with sparse feature masks that allow for direct predic-tion interpretability from the input features. For each forward pass, batch-normalization is applied. The encoder is several steps (parameterized by the user) of self-attention layers and learned sparse feature masks, we offer some preset configurations that depend on the size and complexity of the reference dataset . The decoder then takes these encoded features and passes them through a fully-connected layer with batch-normalization and a generalized linear unit activation [33]. Interpretability by sample is then measured as the sum of feature mask weights across all encoding layers. For our visualiza-tion, we average all feature masks across all cells to understand the average contribution of each gene to the classification. You could also average the feature masks by cell type.

#### 4.1.1. Model Architecture

The encoder architecture consists of three components: a feature trans-former, an attentive transformer, and a feature mask. The raw features are used as inputs, and while no global normalization is applied internally, batch normalization is utilized during training to improve convergence and stabil-ity. [99]. The same *p* dimensional inputs are passed to each decision step of the encoder, which has *N_steps_* decision steps. For feature selection at the *i*th step, an element-wise multiplicative learnable mask *M_i_* is used. This mask is learned via the attentive transformer, and sparsemax normalization [100] is used to induce sparsity in the feature mask. These sequential feature masks are then passed to fully-connected layers for the classification head, first normalized via batch normalization with a gated linear unit [33] for the activation. In our case, we use the raw output of the fully connected classification layer, as [31] loss functions handle logits.

#### 4.1.2. Interpretability

In SIMS the input features correspond to the genes used for cell type prediction by the classifier. Unlike other machine learning models in where computational restrictions force reduced input data representation [101, 41], SIMS can be trained on the entire transcriptome for each cell.

TabNet, which serves as the foundation for SIMS, enables interpretability through the calculation of the weights of the sparse feature masks in the en-coding layer. This allows for an understanding of which input features were utilized in the prediction process at the level of an individual cell. Further-more, by averaging the sum of the attention weights across all samples for a given cell type, it is possible to determine the features used per class, while averaging across all cells in a sample shows the total features used when clas-sifying the entire dataset. Similar to other deep learning models [25], in SIMS the weights do not represent differential gene expression but a measure of the relevance (positive or negative signal) of said gene in the distinction between cell types. Additionally, the sparsity introduced in the sequential attention layers via the sparsemax prior acts as a form of model regularization [30], allowing us to categorize a cell type via only a small number of genes.

### 4.2. Code Library Details

The SIMS pipeline was designed with an easy to use application program-ming interface (API) to support a streamlined analysis with minimal code. To achieve this goal, the pipeline was constructed primarily using PyTorch Lightning, a high-level library that aims to improve reproducibility, modu-larity, and simplicity in PyTorch deep learning code. We utilized Weights and Biases to visualize training metrics, including accuracy, F1 score, and loss, to facilitate the assessment of model performance.

To accommodate the large data formats used by SIMS, we implemented two methods for data loading: a distributed h5 backend for training on h5ad files and a custom parser for csv and delimited files that allows for the incre-mental loading of individual samples during training. These same methods are also used for inference. In addition, cell-type inference can be performed directly on an h5ad file that has been loaded into memory. This allows for efficient handling of datasets that may exceed the available memory capacity. We strongly support the use of h5ad files as they are faster and more efficient than plain text files and allow for more straight forward data sharing in the python-scanpy environment.

All the code and instructions to use SIMS are available in the Braingeneers GitHub repository: https://github.com/braingeneers/SIMS

#### 4.2.1. Web application

In parallel to the API we also developed a Web application in Streamlit. In this case the web application allows for quick and easy inference based on pretrained models. The user only needs to input the single cell RNA dataset in the h5ad format, select the pretrained model they want to use and perform the predictions. The application is hosted in the streamlit developer cloud, allowing access from anywhere without the need of institutional credentials. Laboratories interested in sharing models created with their data with the public can request to include their pretrained models in our repository for easy hosting with a git push request.

#### 4.2.2. Training details

For all models benchmarked, the Adam optimizer [102] was used. The learning rate varied but was generally between 0.003 and 0.01, while the weight decay (L2 regularization) was between 0 and 0.1. To numerically encode the vectors, we used a standard one-hot encoding, where for *K* labels we have that the *k*th label is given by the standard basis vector *e_k_* of all zeros except a 1 in the *k*th position. To define error in the model, average over the categorical cross-entropy loss function, defined as

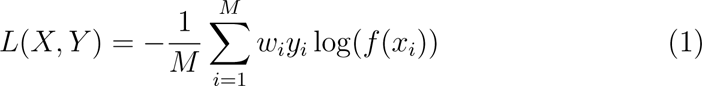

Where *_i_x* represents the transcriptome vector for the *i*th sample, *y_i_* is the encoded label, *w_i_* is the weight and *M* is the size of the batch. For our model, we defined *w_i_* as the inverse frequency of the *i*th label, in order to incentivize the model to learn the transcriptomic structure of rarer cell types. The final signal to update the model weights was calculated as the average across all entries in the loss vector.

A learning rate optimizer was used such that *l ←* 0.75*l* when the vali-dation loss did not improve for twenty epochs. In all cases, models reached convergence by the early stopping criterion on validation accuracy before the maximum number of epochs (500) was reached. Gradient clipping was used to avoid exploding gradient values, which was required to avoid bad batches exploding the loss and stopping convergence. Although we used a train, vali-dation and test split for reducing overfitting via hyperparameter tuning bias, the only hyperparameters tuned were the learning rate to avoid divergence in the loss and weight decay to avoid overfitting in the smaller datasets. Convergence took around 20-100 epochs for all models. For all models, we found model training to be consistent and had few cases of suboptimal convergence due to poor initialization. The train, validation and test sets were stratified, meaning the distribution of labels is the same in all three (up to an error of one sample, when the number of samples for a given class was not divisible by three), except for the ablation study, where there were not enough samples to stratify across all three splits.

For all benchmarks, models were trained using the most granular annotation available. When F1 score is mentioned in benchmarks it refers to the Macro F1-score.

#### 4.2.3. Datasets

##### Peripheral blood mononuclear cells (PBMC68K) dataset

Also known as Zheng68K is the PBMC dataset described in [39]. The dataset was generated using 10X Genomics technologies and sequenced using Illumina NextSeq500. It contains about 68,450 cells within eleven subtypes of cells. The distribution of cell types is imbalanced and transciptomic similarities be-tween cell types makes classification a difficult task. Due to these properties, the PBMC68K dataset is widely used for cell type annotation performance assessment. The dataset can be accessed at https://www.10xgenomics.com/resources/datasets/fresh-68-k-pbm-cs-donor-a-1-standard-1-1-0

##### Human cellular landscape: Han’s dataset

The Human cellular landscape dataset described in [103]. The dataset was generated using Microwell-seq technology. It contains 584000 cells with 102 different cell types across all major human organs and different developmental timepoints from more than 50 different donors. The data can be accesed at https://cells.ucsc.edu/?ds=human-cellular-landscape

##### Human Heart: Tucker’s dataset

The Tucker dataset described in [40] is a single nuclei RNA-sequencing dataset comprised of 287,269 cells representing 9 different cell types (20 cell subtypes) from 7 different donors. The dataset was acquired from https://singlecell.broadinstitute.or g/single_cell/study/SCP498/transcriptional-and-cellular-diver sity-of-the-human-heart#study-summary

##### Adult mouse cortical and hippocampal dataset

This dataset was generated by the Allen Brain Institute and described in [43, 44, 45]. The dataset was generated from male and female 8 week-old mice labeled using pan-neuronal transgenic lines. The dataset includes micro-dissected corti-cal and hippocampal regions. It contains 42 cell types including excitatory projection neurons, interneurons and non-neuronal cells. The dataset can be accessed at https://portal.brain-map.org/atlases-and-data/rnaseq/mouse-whole-cortex-and-hippocampus-10x

##### Adult human cortical dataset

This dataset was generated from post-mortum samples by the Allen Brain Institute [44, 43]. It includes single-nucleus transcriptomes from 49,495 nuclei across multiple human cortical areas. The large majority of nuclei are contributed from 3 donors: 1) H200-1023 was a female Iranian-descent donor who was 43 years old at the time of death. The cause of death was mitral valve collapse. 2) H200-1025 was a male Caucasian donor who was 50 years old at the time of death. The cause of death was cardiovascular. 3) H200-1030 was a male Caucasian donor who was 57 years old at the time of death. The cause of death was cardiovascu-lar. For sampling, individual cortical layers were dissected from the middle temporal gyrus, anterior cingulate cortex, primary visual cortex, primary motor cortex, primary somatosensory cortex and primary auditory cortex. All samples were dissected from the left hemisphere. As part of the purifica-tion processes, nuclei were isolated and sorted using Fluorescently Activated Cell Sorting (FACS) using NeuN as a marker. For statistics, we only used cell types that were common between all samples. The data was obtained from https://portal.brain-map.org/atlases-and-data/rnaseq/human-multiple-cortical-areas-smart-seq.

##### Developing mouse cortical dataset

This dataset was described in [60]. It contains microdissected cortices from mice ranging from embryonic day 10 to postnatal day 4. For this study we used data from mice at em-bryonic day 12 (1 batch, 9,348 cells), 13 (1 batch, 8,907 cells), 14 (1 batch, 5249 cells) and 18 (2 batches, 7,137 cells), as well as postnatal day 1 (2 batches, 13,072 cells). Of note, only postnatal day 1 samples had Ependy-mocytes, and as such, they were removed for inter-age testing. The data was downloaded from the Single Cell Portal administered by the Broad Institute. https://singlecell.broadinstitute.org/single_cell/study/SCP129 0/molecular-logic-of-cellular-diversification-in-the-mammalian-cerebral-cortex

##### Human cortical organoids dataset

We used 6-months old organoids described in [74]. The dataset contained cells derived from 3 cell lines: GM8330 (3 organoids, 1 batch, 15,256 cells), 11A (3 organoids, 1 batch, 25,618 cells) and PGP1 (6 organoids 2 batches, 46,989 cells). PGP1 has a strong batch effect which is almost entirely caused by one organoid in batch 3. The dataset was generated using Chromium Single Cell 3’ Library and Gel Bead Kit v.2 (10x Genomics, PN-120237) and sequenced using the Illumina NextSeq 500 instrument. Of note, one of the cell lines had a cell cluster named “Callosal Projection Neurons” while others had “Immature Callosal Projection Neurons. Given the naming inconsistency, we aggregated both clusters as “Callosal Projection Neurons”. We downloaded the dataset from the Single Cell Portal administered by the Broad Institute. https://singlecell.broadinstitute.org/single_cell/study/SCP282/reproducible-brain-organoids#study-summary

##### Human fetal brain development

We utilized fetal tissue representative of the second trimester of human development, specifically focusing our analysis on data sourced exclusively from the neocortex. This study encompassed the sampling of six distinct neocortical regions. The dataset contained samples from gestational weeks 14, 17, 18, 19, 20, 22, and 25. The number of cells contained in this dataset was around 404000 [83]. https://cells.ucsc.edu/?bp=brain&ds=dev-brain-regions

### 4.3. Benchmarking against cell type classification models

We benchmarked SIMS using the Zheng68K and Tucker’s dataset, as previously described[25].We also added Han’s dataset to the benchmark. Briefly, we compared our model to:

#### scBERT 1.0

scBERT is a transformer architecture based on the deep learning model BERT. It has been adapted to work with single cell data and it offers interpretability as the attention weights for each gene. [25]

#### scNym 0.3.2

scNym is a neural network model for predicting cell types from single cell profiling data and deriving cell type representations from these models. These models can map single cell profiles to arbitrary output classes. [28]

#### scANVI 1.0.2

scANVI (single-cell ANnotation using Variational Inference) represents a semi-supervised approach designed specifically for single-cell transcriptomics data. It relies on the utilization of variational autoen-coders as the foundational component of its model architecture[27]

#### SciBet 1.0

SciBet is a supervised classification tool, consisting of 4 steps: preprocessing, feature selection, model training and cell type assignment, that selects genes using E-test for multinomial model building. [41]

#### Seurat 4.0.3

We used Seurat’s reference-based mapping, with the Transfer anchor settings, where very transcriptomically simmilar cells from the reference and query datasets are used to create a shared space for the two datasets[19]

#### SingleR 1.6.1

SingleR is a reference-based method that requires tran-scriptomic datasets of pure cell types to infer the cell of origin of each of the single cells independently. It uses the Spearman coefficient on variable genes and aggregates the coefficients to score the cell for each cell type[20]

### 4.4. Pseudotime analysis: Monocle 3.1

The human cortical organoid dataset was parsed into R (v. 4.2.1) using Seurat and its dependencies (v. 4.3.0) and converted into a CellDataSet (CDS) for further analysis using Monocle 3 Beta (v. 3.1.2.9; https://cole-trapnell-lab.github.io/monocle3/) [84]. Cell clusters and trajectories were visualized utilizing the conventional Monocle workflow, as detailed in https://cole-trapnell-lab.github.io/monocle3/docs/trajectories/.

### 4.5. Cell stress analysis: Gruffi 1.0

Gruffi is a computational algorithm that identifies and removes stressed cells from brain organoid transcriptomic datasets in an unbiased manner [81]. It uses granular functional filtering to isolate stressed cells based on stress pathway activity scoring [81]. Gruffi integrates into a typical single-cell analysis workflow using Seurat [81]. In this paper we followed the default implementation shown in the GitHub repository to obtain a dataframe con-taining what cells were stressed based on Gruffi’s default analysis https://github.com/jn-goe/gruffi.

## 5. Declarations

### 5.1. Author Contribution Statement

B.P., M.T., D.H., V.D.J., and M.A.M.-R. conceived the project. J.G.-F. and J.L. performed the experiments. A.O. provided support working with the Terra system. J.G.-F. J.L., and M.A.M-R. wrote the paper with contributions from all authors.

### 5.2. Data Availability Statement

All data used in this paper comes from previously published datasets. Peripheral blood mononuclear cells: https://www.10xgenomics.com/resources/datasets/fresh-68-k-pbm-cs-donor-a-1-standard-1-1-0

Human cellular landscape: https://cells.ucsc.edu/?ds=human-cellular-landscape

Tucker’s heart dataset: https://singlecell.broadinstitute.org/single_cell/study/SCP498/transcriptional-and-cellular-diversity-of-the-human-heart

Human adult cerebral cortex: https://portal.brain-map.org/atlases-and-data/rnaseq/human-multiple-cortical-areas-smart-seq

Mouse adult cerebral cortex and hippocampus: https://portal.brain-map.org/atlases-and-data/rnaseq/mouse-whole-cortex-and-hippocampus-10x

Developing mouse cerebral cortex (E12-P1): https://singlecell.broadinstitute.org/single_cell/study/SCP1290/molecular-logic-of-cellular-diversification-in-the-mammalian-cerebral-cortex

Human cortical organoids: https://singlecell.broadinstitute.org/single_cell/study/SCP282/reproducible-brain-organoids#study-summary

Human fetal brain development: https://cells.ucsc.edu/?bp=brain&ds=dev-brain-regions

## 5.4 Acknowledgments

We would like to thank Tomasz Nowakowski, Maximilian Haeussler, Bene-dict Paten and Hunter Schweiger for their valuable feedback on this manuscript. This work was supported by Schmidt Futures (SF857) to M.T. and D.H.; National Human Genome Research Institute (1RM1HG011543) to M.T. and D.H.; National Science Foundation (NSF2134955) to M.T. and D.H. (NSF2034037) to M.T.; the National Institute of Mental Health (1U24MH132628) to B.P., D.H. and M.A.M.-R. We are thankful to the Pacific Research Platform, sup-ported by the National Science Foundation under Award Numbers CNS-1730158, ACI-1540112, ACI-1541349, OAC-1826967, the University of Cali-fornia Office of the President, and the University of California San Diego’s California Institute for Telecommunications and Information Technology/Qualcomm Institute.

