## Supplemental Materials for "Unraveling Neuronal Identities Using SIMS: A Deep Learning Label Transfer Tool for Single-Cell RNA Sequencing Analysis"

**Figures**:

Supplemental Figure 1: SIMS pipeline
Supplemental Figure 2: SIMS training snippets
Supplemental Figure 3: SIMS predicting snippets
Supplemental Figure 4: Terra workflow
Supplemental Figure 5: Benchmarking of SIMS against common cell labeling algorithms.
Supplemental Figure 6: Training data for the adult mouse cerebral cortex and hippocampus model.
Supplemental Figure 7: Ablation studies in adult mouse cerebral cortex and hippocampus.
Supplemental Figure 8: Explainability of PVALB+ Interneurons in adult human cerebral cortex label transfer, medium granularity.
Supplemental Figure 9: Training data for the adult human cerebral cortex model.
Supplemental Figure 10: Explainability for 300 runs, median genes
Supplemental Figure 11: Ablation studies in adult human cerebral cortex.
Supplemental Figure 12: Trans-sample label transfer in adult human cerebral cortex.
Supplemental Figure 13: Label transfer in E18 and P1 developing mouse cerebral cortex.
Supplemental Figure 14: Batch effects in single-cell RNA sequencing of 6 months old PGP1-derived cortical organoids.
Supplemental Figure 15: Heatmap Intra cell line label transfer, 11A
Supplemental Figure 16: Wilcoxon test ranks for the different cell types in the cortical organoids
Supplemental Figure 17: Label transfer from Fetal tissue
Supplemental Figure 18: UMAPs for PGP1 and GM8330
Supplemental Figure 19: UMAP representation with Markers for 11A organoids

**Tables**

Supplemental Table 1: Accuracies of single cell classifiers in the Benchmark datasets.
Supplemental Table 2: Macro F1-scores of single cell classifiers in the Benchmark dataset.
Supplemental Table 3: Accuracies by cell type for sample H200.1025 predictions training SIMS on Sample H200.1023.
Supplemental Table 4: Accuracies by cell type for sample H200.1030 predictions training SIMS on Sample H200.1023.

Supplemental Table 5: Accuracies by cell type for sample H200.1023 predictions training SIMS on Sample H200.1025.
Supplemental Table 6: Accuracies by cell type for sample H200.1030 predictions training SIMS on Sample H200.1025.
Supplemental Table 7: Accuracies by cell type for sample H200.1023 predictions training SIMS on Sample H200.1030.
Supplemental Table 8: Accuracies by cell type for sample H200.1025 predictions training SIMS on Sample H200.1030.
Supplemental Table 9: Accuracies by cell type for E18 and P1 samples predictions training SIMS on E18 and P1 samples.
Supplemental Table 10: Accuracies by cell type for E18 samples predictions training SIMS on P1 samples.
Supplemental Table 11: Accuracies by cell type for GM8330-derived organoids predictions training SIMS on 11A-derived organoids.
Supplemental Table 12: Accuracies by cell type for PGP1-derived organoids predictions training SIMS on 11A-derived organoids.
Supplemental Table 13: Accuracies by cell type for GM8330-derived organoids predictions training SIMS on 11A-derived organoids after reclassification of 11A cells.
Supplemental Table 14: Accuracies by cell type for PGP1-derived organoids predictions training SIMS on 11A-derived organoids

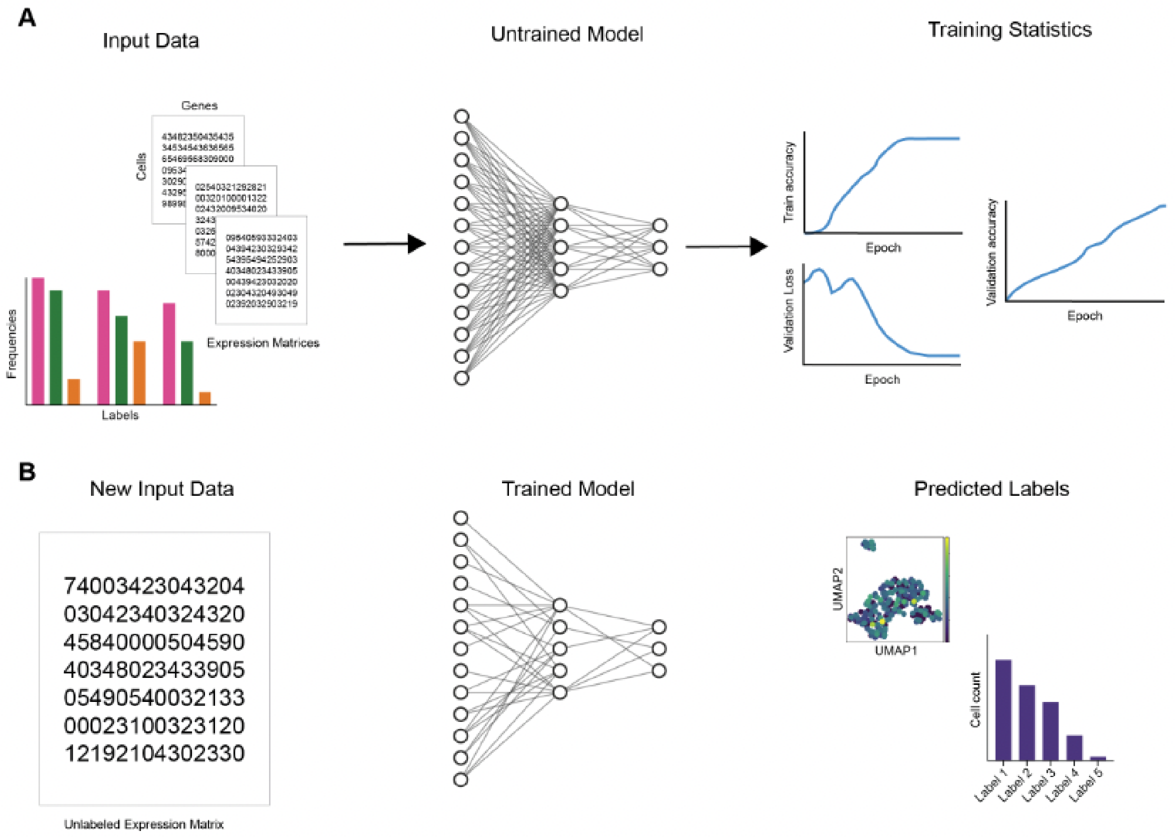


**Supplemental Figure 1. SIMS pipeline.** A) Training step, input data in the form of a cell by gene matrix is fed to the algorithm to train it. The training statistics can be followed live. B) Prediction step, input data is fed to the algorithm which outputs labels and a confidence score.

```python
from scsims import SIMS

# set up the SIMS model to train on a column in the .obs
sims = SIMS(
    data="my_data.h5ad",
    class_label='cell_type',
)

# train the model
sims.train(max_epochs=100)
```

**Supplemental Figure 2. SIMS training snippet**

```python
from scsims import SIMS

# set up the SIMS model to run inference
# from a model saved during training
sims = SIMS(
    weights_path="my_model.ckpt",
)

# get the predicted cell types and probabilities
predictions = sims.predict("unlabeled_data.h5ad")
```

**Supplemental Figure 3. SIMS prediction snippet**

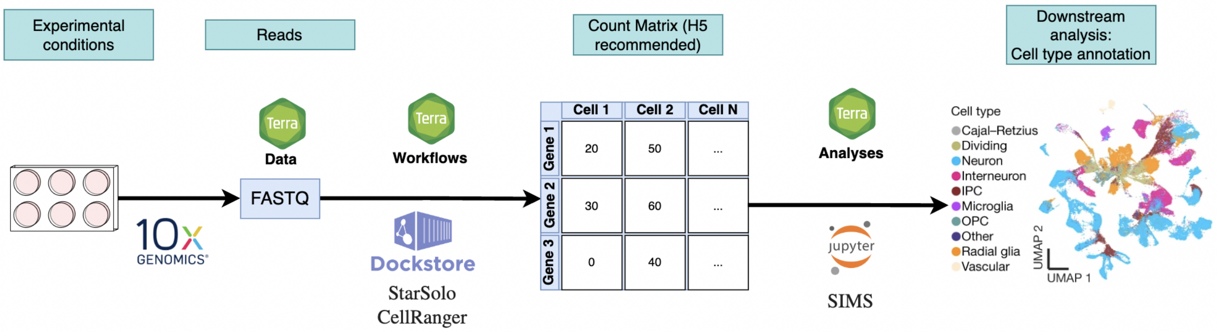


**Supplemental Figure 4. Suggested Terra Workflow.** This workflow takes raw data from single cell experiments in the FASTQ format, runs them through dockstore workflows for data integration and count matrix creation and performs analysis in the Terra notebooks workspace.

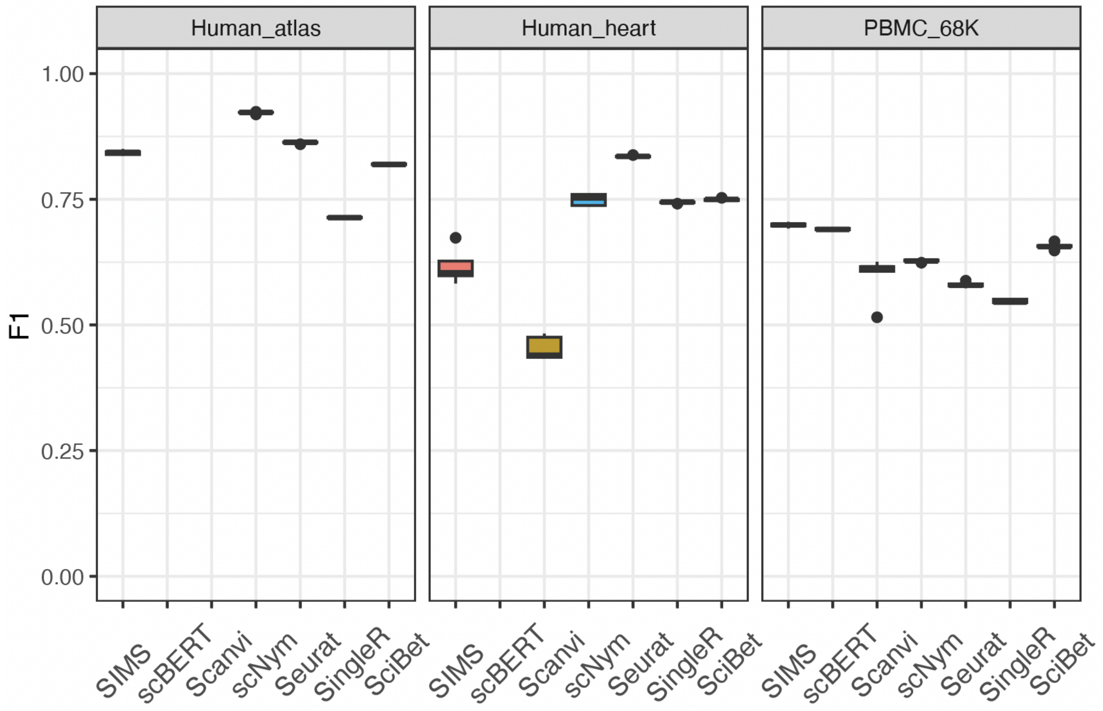


**Supplemental Figure 5. Benchmarking of SIMS against common cell labeling algorithms.** Macro F1 scores of SIMS and other models using the Human Cell Atlas, Human Heart and PBMC68K dataset.

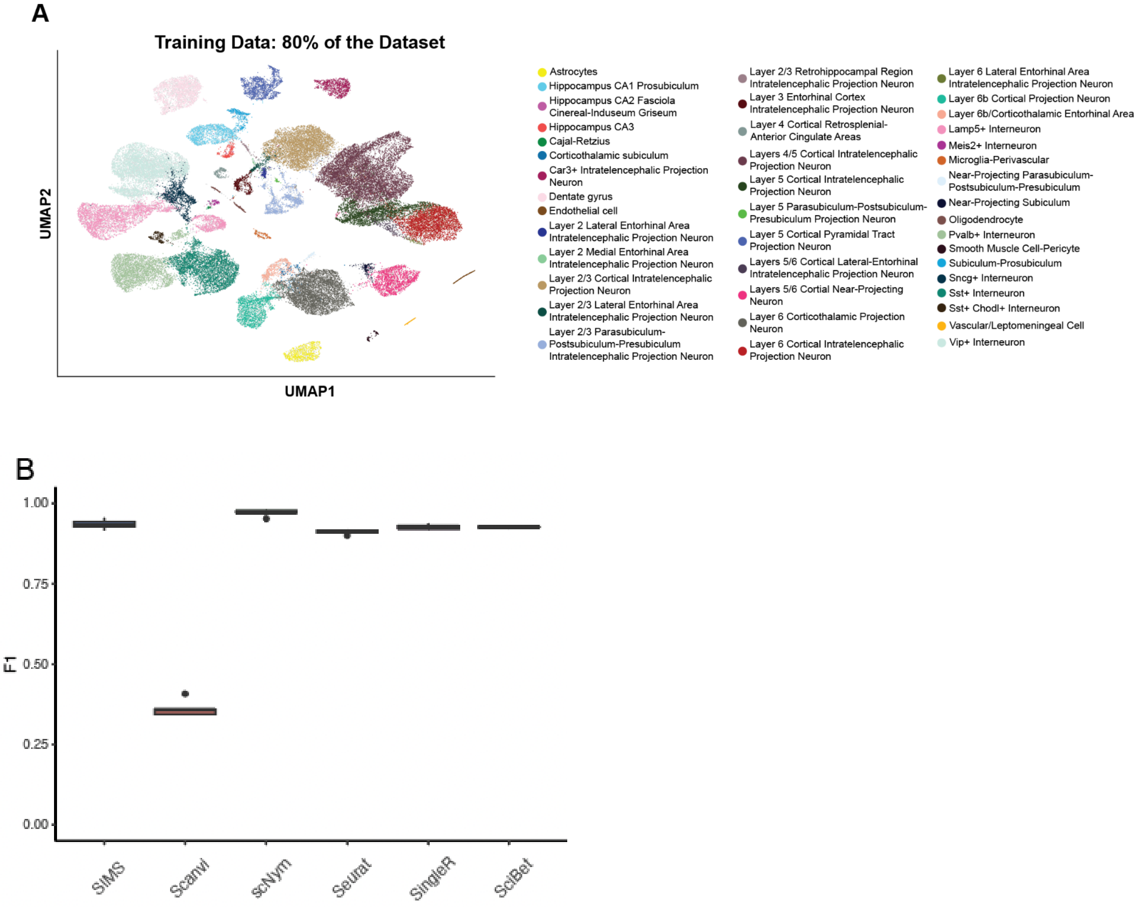


**Supplemental Figure 6. Training data for the adult mouse cerebral cortex and hippocampus model.** A) UMAP of the training data for the adult mouse cerebral cortex and hippocampus label transfer experiment. Cells non used for training are shown in light gray. B) F1 score for Allen mouse benchmark

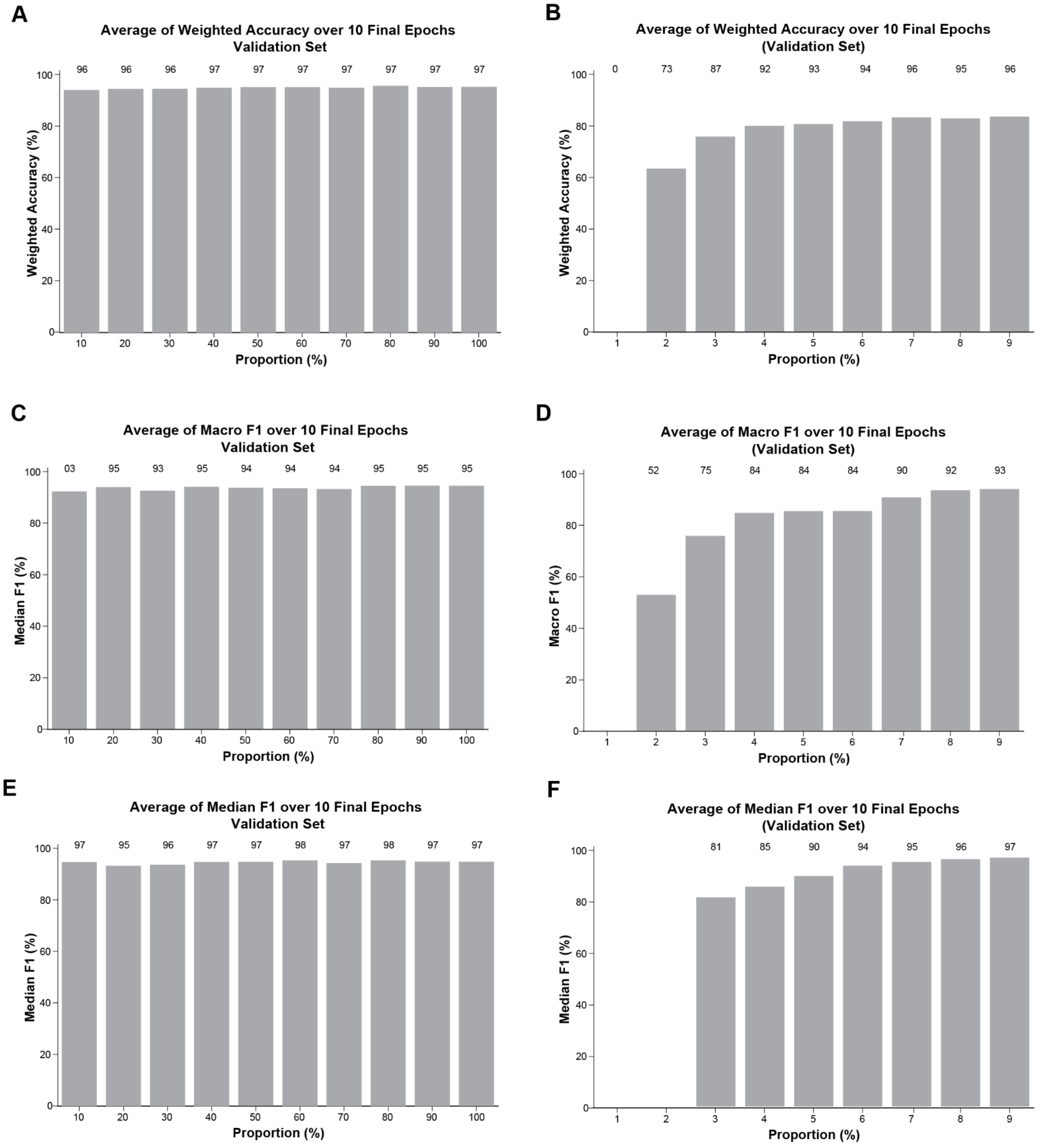


**Supplemental Figure 7. Ablation studies in adult mouse cerebral cortex and hippocampus.** A) Average of Weighted Accuracy over 10 Final Epochs (10-100%). B) Average of Weighted Accuracy over 10 Final Epochs (1-9%). C) Average of Macro F1 over 10 Final Epochs (10-100%). D) Average of Macro F1 over 10 Final Epochs (1-9%). E) Average of Median F1 over 10 Final Epochs (10-100%). F) Average of Median F1 over 10 Final Epochs (1-9%).

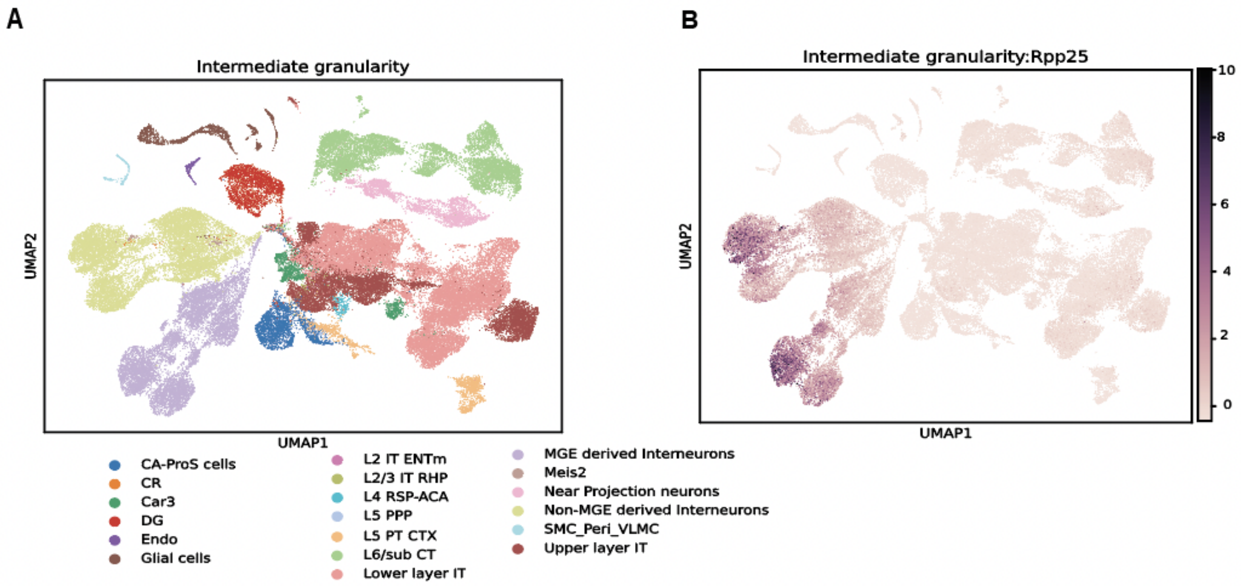


**Supplemental Figure 8: Explainability of PVALB+ Interneurons in adult human cerebral cortex label transfer, Medium granularity.**

A

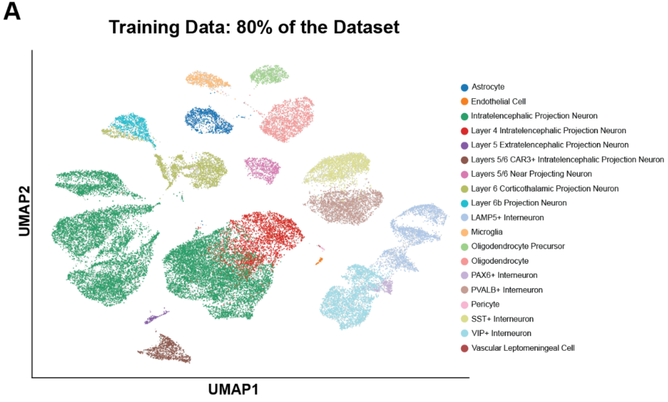


**Supplemental Figure 9. Training data for the adult human cerebral cortex model.** A) UMAP of the training data for the adult human cerebral cortex label transfer experiment. Cells non used for training are shown in light gray.

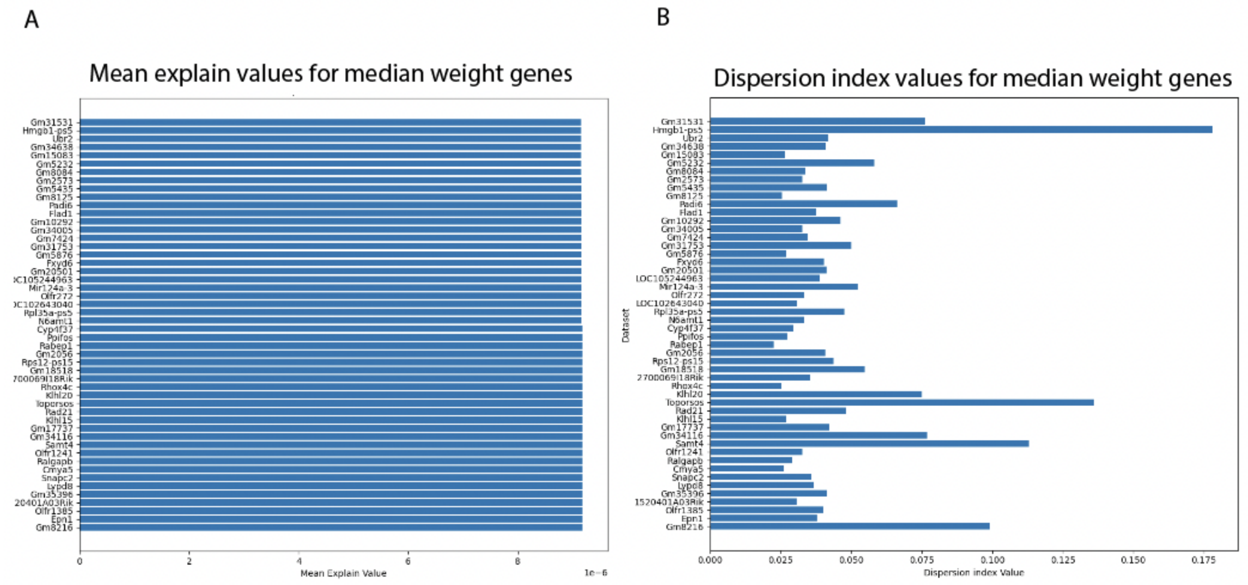


**Supplemental Figure 10. Explainability for 300 runs, median genes.** A) Mean explain values for median genes. B) Dispersion index for median genes

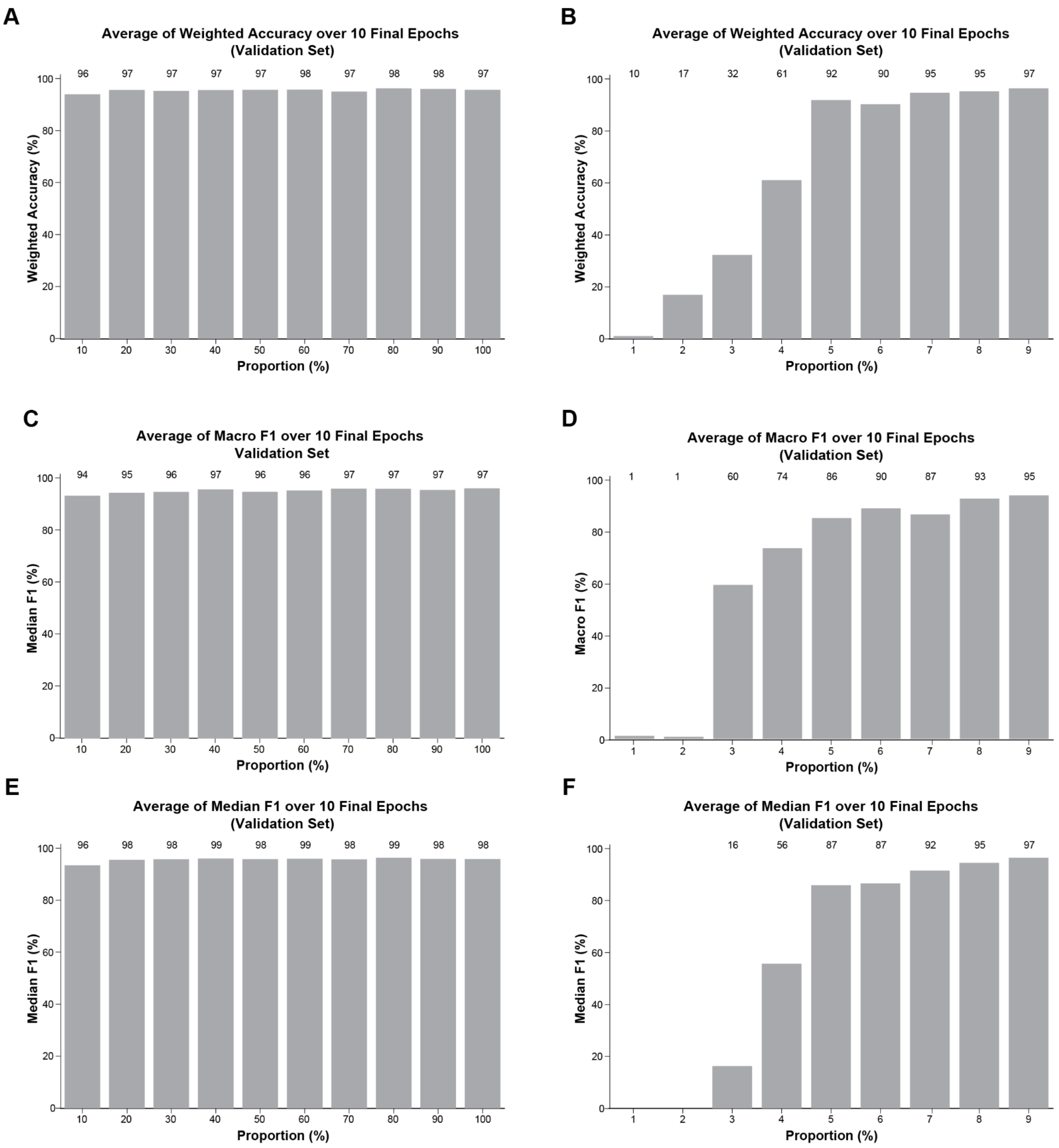


**Supplemental Figure 11. Ablation studies in adult human cerebral cortex.** A) Average of Weighted Accuracy over 10 Final Epochs (10-100%). B) Average of Weighted Accuracy over 10 Final Epochs (1-9%). C) Average of Macro F1 over 10 Final Epochs (10-100%). D) Average of Macro F1 over 10 Final Epochs (1-9%). C) Average of Median F1 over 10 Final Epochs (10-100%). D) Average of Median F1 over 10 Final Epochs (1-9%).

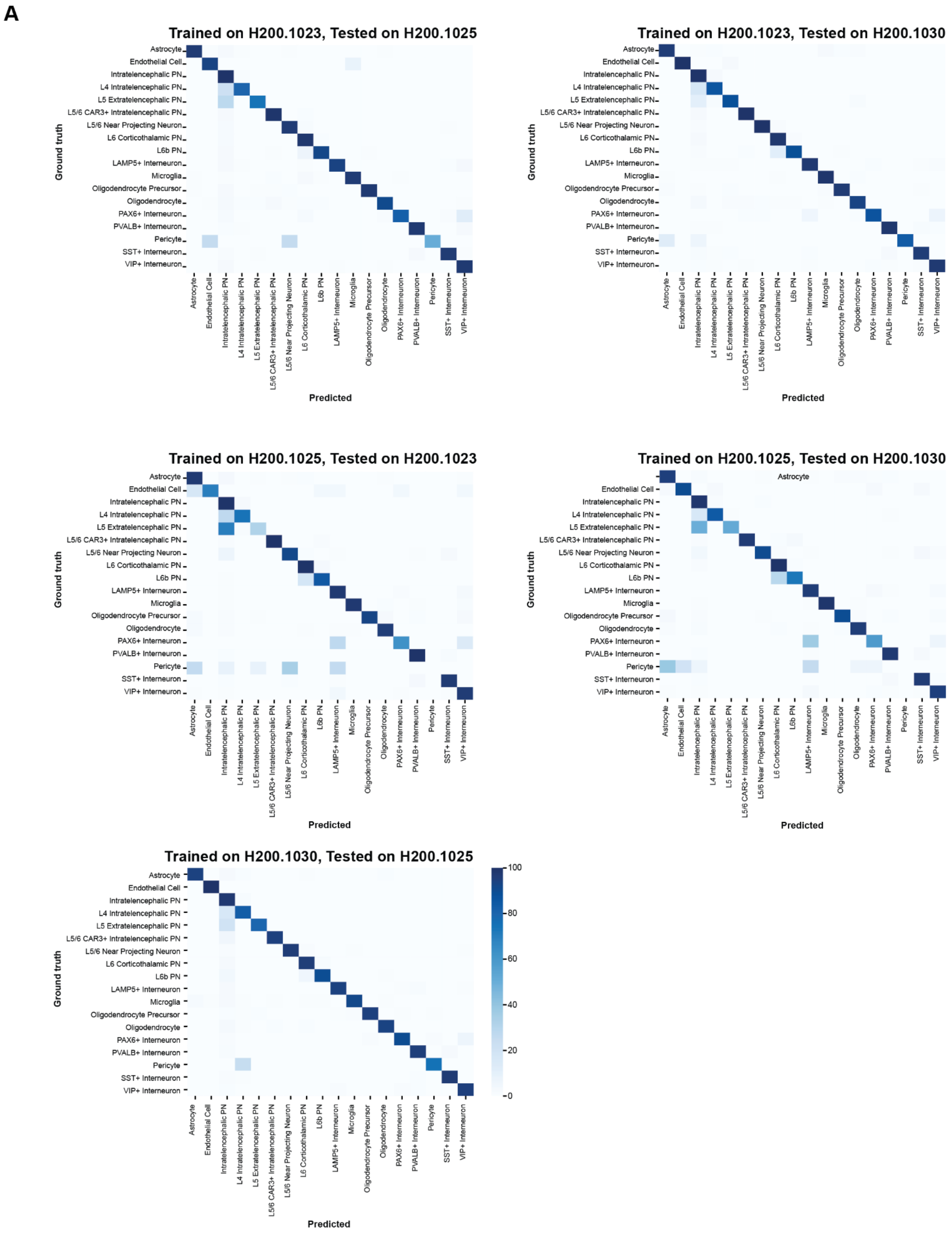


**Supplemental Figure 12. Trans-sample label transfer in adult human cerebral cortex.** A) Remaining combinations of training and testing label transfer across 3 samples of the adult human cerebral cortex: H200.1023, H200.1025 and H200.1030.

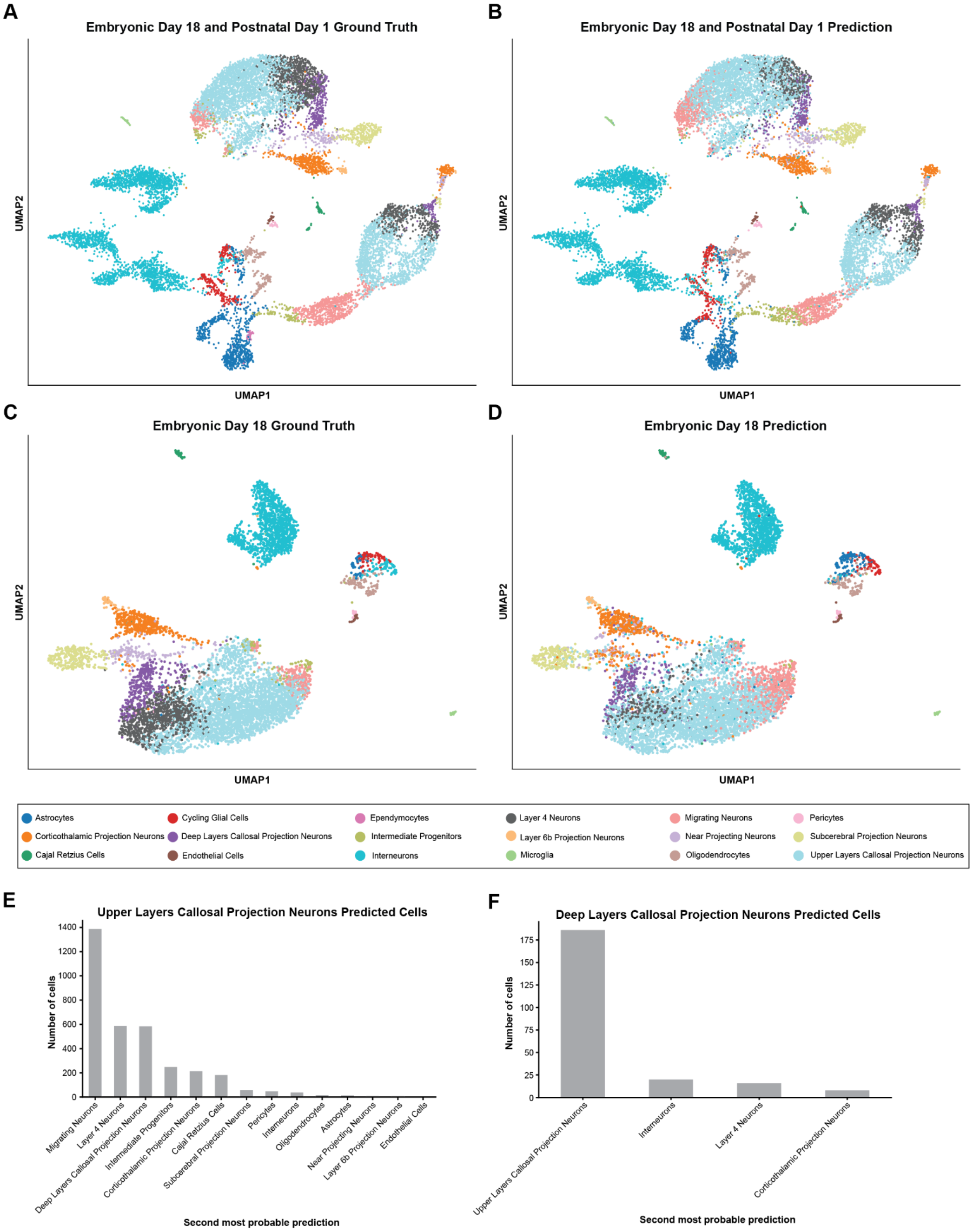


**Supplemental Figure 13. Label transfer in E18 and P1 developing mouse cerebral cortex.** A) UMAP representing the ground truth of cell types present in one sample of E18 and one sample of P1 mouse cerebral cortex. B) Prediction of the same E18 and P1 samples after training SIMS in different E18 and P1 samples. C) UMAP representing the ground truth of cell types present in two samples of E18 mouse cerebral cortex. D) Prediction of the same E18 samples after training SIMS in two P1 samples. E) Second most probable prediction for upper layer callosal projection neurons. F) Second most probable prediction for deep layer callosal projection neurons.

A

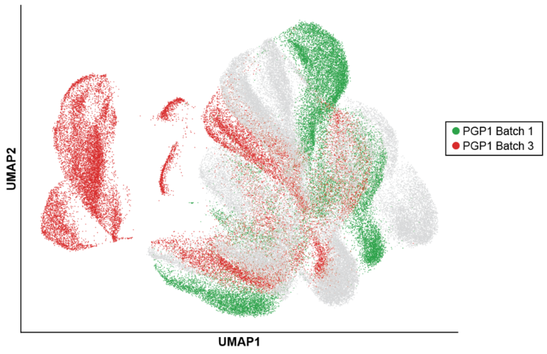


**Supplemental Figure 14. Batch effects in single-cell RNA sequencing of 6 months old PGP1-derived cortical organoids.** A) UMAP representation of cells from PGP1-derived cortical organoids. Green = Cells from PGP1 batch 1. Red = Cells from PGP1 batch 3. Gray = Cells from GM8330 and 11A-derived cortical organoids.

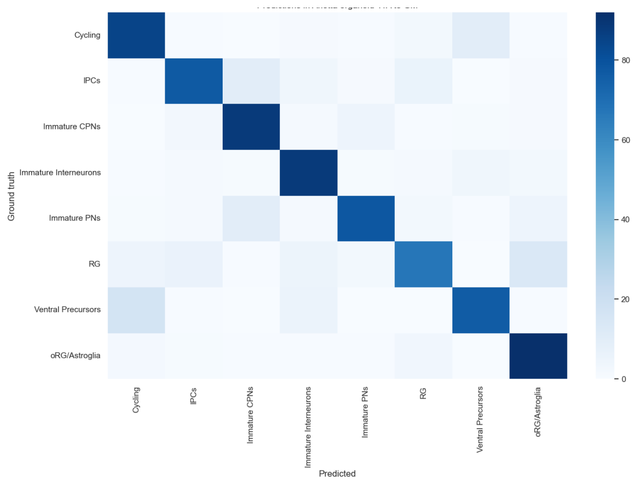


**Supplemental Figure 15**: **Heatmap Intra cell line label transfer**. Heatmap for the 11A cell line intra c. We trained the algorithm in two 11A organoids and predicted on a third 11A organoid

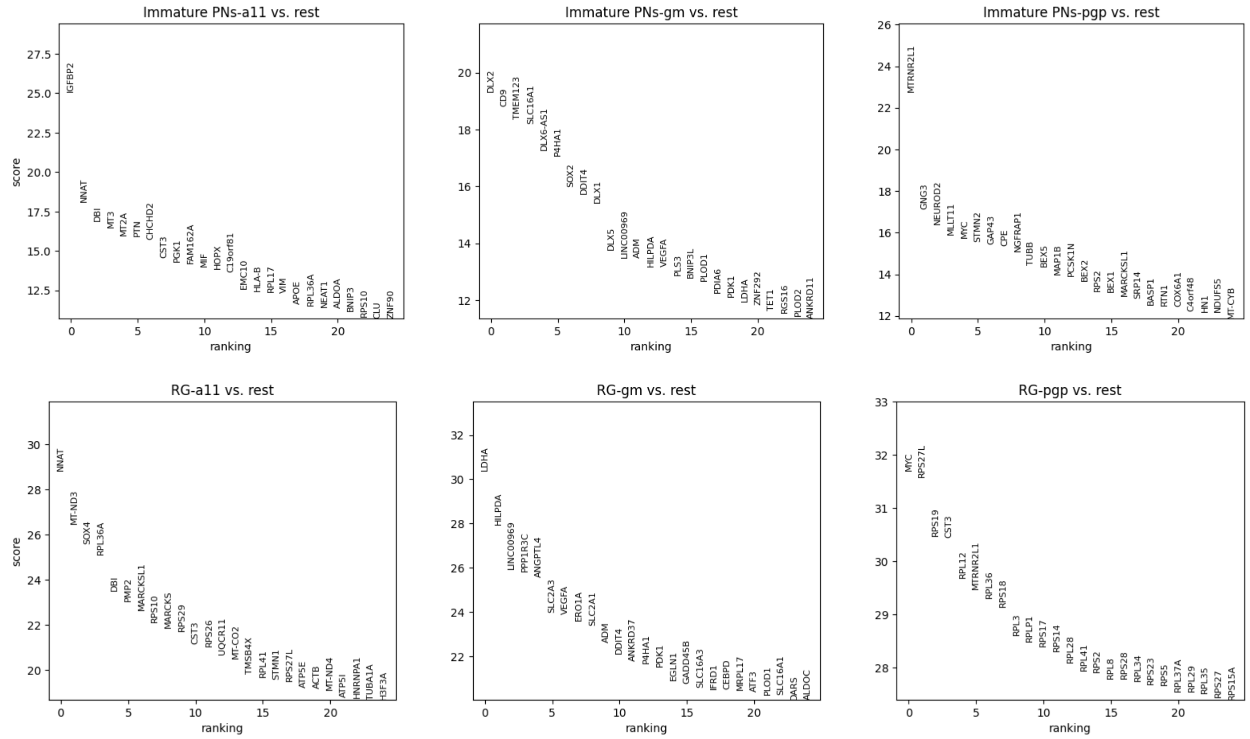


**Supplemental Figure 16**: **Wilcoxon test ranks for Radial Glia and Immature PNs in the cortical organoids**

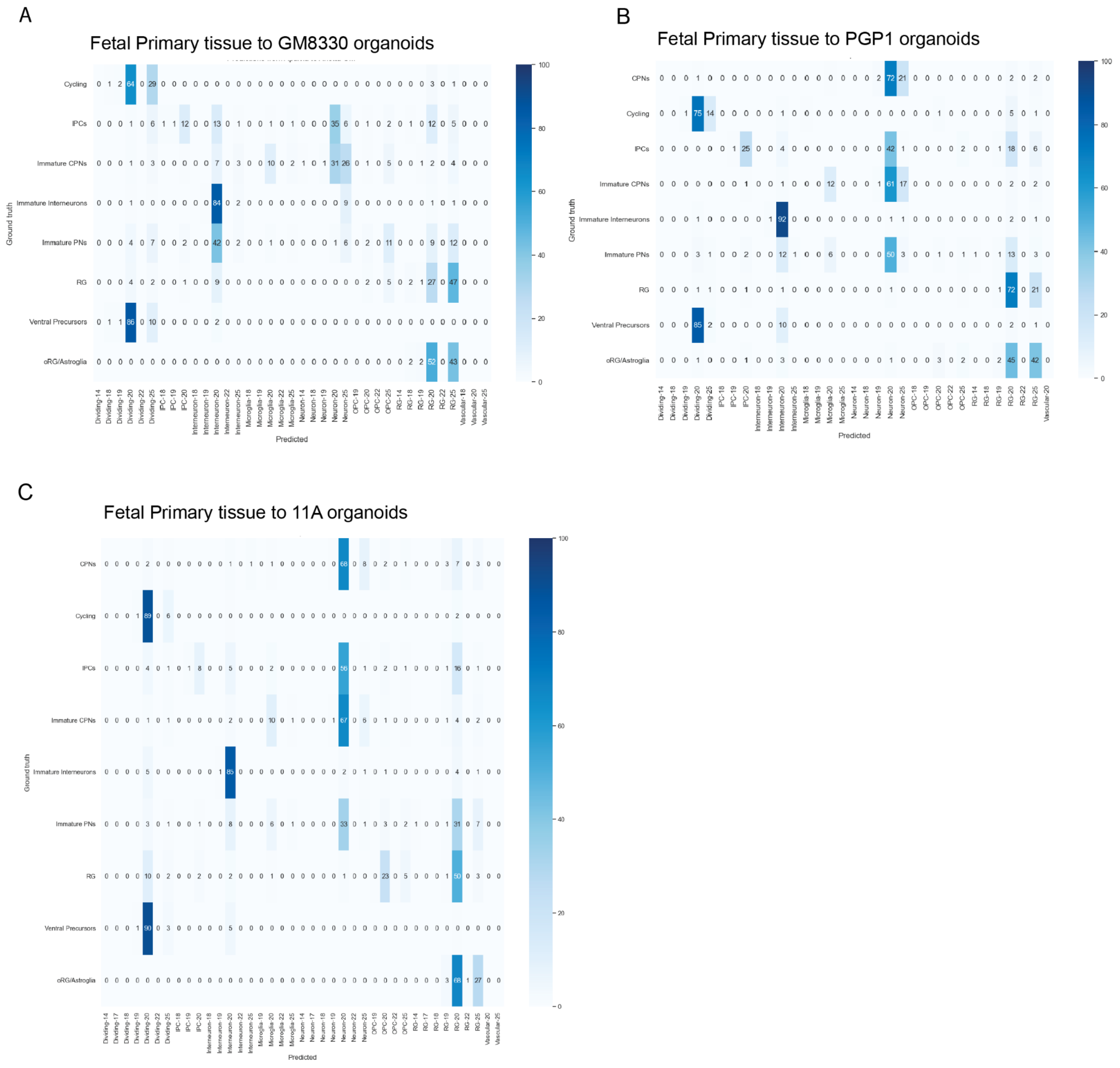


**Supplemental Figure 17: Heatmaps showing Label transfer from Fetal tissue to organoids from different cell lines**

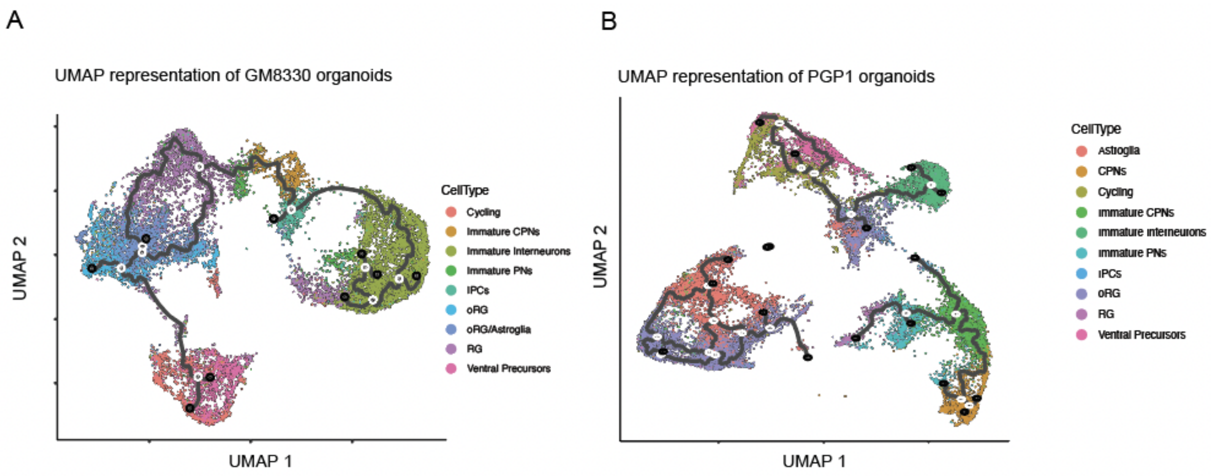


**Supplemental Figure 18: UMAPS for PGP1 and GM8330.**

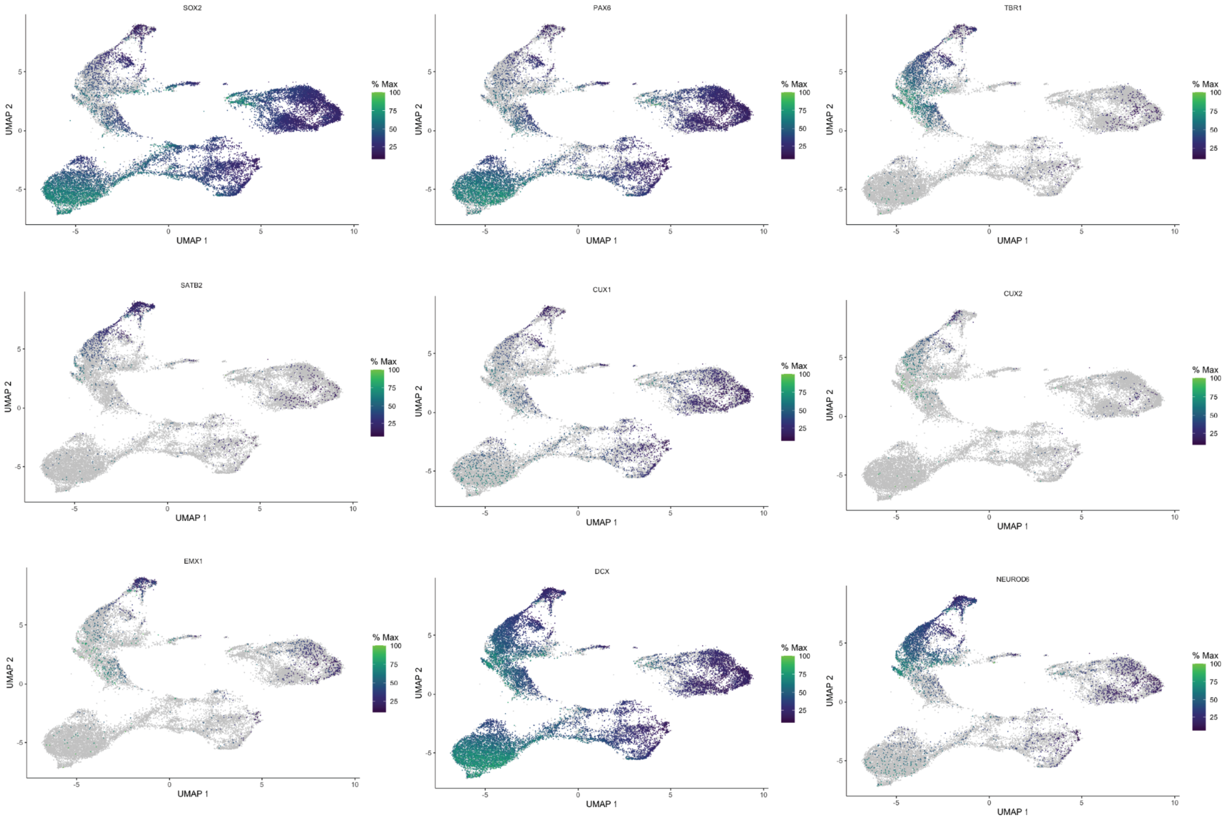


**Supplemental Figure 19: UMAP representation with Markers for 11A organoids**

**Supplemental Table 1. Accuracies of single cell classifiers in the Benchmark datasets**

| Measure Accuracy | Dataset | Tool |
|---|---|---|
| 0.890741343 | Human_atlas | SIMS |
| 0.8917598124 | Human_atlas | SIMS |
| 0.8890210712 | Human_atlas | SIMS |
| 0.887959809 | Human_atlas | SIMS |
| 0.8893377381 | Human_atlas | SIMS |
| 0.9264592605 | Human_atlas | scNym |
| 0.9255178021 | Human_atlas | scNym |
| 0.9271225608 | Human_atlas | scNym |
| 0.9257103732 | Human_atlas | scNym |
| 0.9286631291 | Human_atlas | scNym |
| 0.8315588573 | Human_atlas | SciBet |
| 0.8301980452 | Human_atlas | SciBet |
| 0.832431831 | Human_atlas | SciBet |
| 0.831464713 | Human_atlas | SciBet |
| 0.8792547137 | Human_atlas | Seurat |
| 0.879246155 | Human_atlas | Seurat |
| 0.8779709177 | Human_atlas | Seurat |
| 0.8768754119 | Human_atlas | Seurat |
| 0.8765245077 | Human_atlas | Seurat |
| 0.7188938909 | Human_atlas | SingleR |
| 0.719167765 | Human_atlas | SingleR |
| 0.7186200168 | Human_atlas | SingleR |
| 0.7176956916 | Human_atlas | SingleR |
| 0.7696593011 | Human_heart | SIMS |
| 0.7577691396 | Human_heart | SIMS |
| 0.7534246575 | Human_heart | SIMS |
| 0.7595568213 | Human_heart | SIMS |
| 0.8111293575 | Human_heart | SIMS |
| 0.8432157148 | Human_heart | scNym |
| 0.8447227563 | Human_heart | scNym |
| 0.8454502936 | Human_heart | scNym |
| 0.8484643767 | Human_heart | scNym |
| 0.8419165411 | Human_heart | scNym |

| | | |
|---|---|---|
| **0.7711767518** | Human_heart | Scanvi |
| **0.7620512607** | Human_heart | Scanvi |
| **0.7667075478** | Human_heart | Scanvi |
| **0.7864967676** | Human_heart | Scanvi |
| **0.7775583595** | Human_heart | Scanvi |
| **0.772357016** | Human_heart | SciBet |
| **0.772357016** | Human_heart | SciBet |
| **0.7692066697** | Human_heart | SciBet |
| **0.7693111011** | Human_heart | SciBet |
| **0.7696940161** | Human_heart | SciBet |
| **0.8449863367** | Human_heart | Seurat |
| **0.8447948758** | Human_heart | Seurat |
| **0.8452648252** | Human_heart | Seurat |
| **0.8431413503** | Human_heart | Seurat |
| **0.8477712217** | Human_heart | Seurat |
| **0.7434991471** | Human_heart | SingleR |
| **0.7439168726** | Human_heart | SingleR |
| **0.7455007484** | Human_heart | SingleR |
| **0.745622585** | Human_heart | SingleR |
| **0.7436209837** | Human_heart | SingleR |
| **0.7402** | PBMC_68K | SIMS |
| **0.7546** | PBMC_68K | SIMS |
| **0.7548** | PBMC_68K | SIMS |
| **0.7466** | PBMC_68K | SIMS |
| **0.7503** | PBMC_68K | SIMS |
| **0.76** | PBMC_68K | scBERT |
| **0.755** | PBMC_68K | scBERT |
| **0.754** | PBMC_68K | scBERT |
| **0.752** | PBMC_68K | scBERT |
| **0.774** | PBMC_68K | scBERT |
| **0.694521** | PBMC_68K | scNym |
| **0.701241** | PBMC_68K | scNym |
| **0.713075** | PBMC_68K | scNym |
| **0.694302** | PBMC_68K | scNym |
| **0.69853** | PBMC_68K | scNym |
| **0.75386410** | PBMC_68K | Scanvi |

| | | |
|---|---|---|
| **0.7621755** | PBMC_68K | Scanvi |
| **0.7731116943715370** | PBMC_68K | Scanvi |
| **0.75313502** | PBMC_68K | Scanvi |
| **0.76494604** | PBMC_68K | Scanvi |
| **0.6775** | PBMC_68K | SciBet |
| **0.6809** | PBMC_68K | SciBet |
| **0.6757** | PBMC_68K | SciBet |
| **0.6741** | PBMC_68K | SciBet |
| **0.686512** | PBMC_68K | SciBet |
| **0.6839** | PBMC_68K | Seurat |
| **0.6883** | PBMC_68K | Seurat |
| **0.6862** | PBMC_68K | Seurat |
| **0.681** | PBMC_68K | Seurat |
| **0.6919** | PBMC_68K | Seurat |
| **0.3173** | PBMC_68K | SingleR |
| **0.3267** | PBMC_68K | SingleR |
| **0.324** | PBMC_68K | SingleR |
| **0.3246** | PBMC_68K | SingleR |
| **0.337** | PBMC_68K | SingleR |

### Supplemental Table 2. Macro F1-scores of single cell classifiers in the Benchmark datasets

| F1 | Dataset | Tool |
|---|---|---|
| 0.8447592253 | Human_atlas | SIMS |
| 0.8499620517 | Human_atlas | SIMS |
| 0.8433636062 | Human_atlas | SIMS |
| 0.8369518913 | Human_atlas | SIMS |
| 0.8390889883 | Human_atlas | SIMS |
| 0.9228504023 | Human_atlas | scNym |
| 0.9227343304 | Human_atlas | scNym |
| 0.9240401235 | Human_atlas | scNym |
| 0.9188093776 | Human_atlas | scNym |
| 0.9230278601 | Human_atlas | scNym |
| 0.8188885933 | Human_atlas | SciBet |
| 0.8186075424 | Human_atlas | SciBet |
| 0.8203570627 | Human_atlas | SciBet |
| 0.8199298475 | Human_atlas | SciBet |
| 0.8627578352 | Human_atlas | Seurat |
| 0.8638023345 | Human_atlas | Seurat |
| 0.8638803849 | Human_atlas | Seurat |
| 0.8634841494 | Human_atlas | Seurat |
| 0.8594598098 | Human_atlas | Seurat |
| 0.7137920926 | Human_atlas | SingleR |
| 0.7157409671 | Human_atlas | SingleR |
| 0.7125192737 | Human_atlas | SingleR |
| 0.7132709792 | Human_atlas | SingleR |
| 0.6270632609 | Human_heart | SIMS |
| 0.5826348894 | Human_heart | SIMS |
| 0.6036579798 | Human_heart | SIMS |
| 0.5978908132 | Human_heart | SIMS |
| 0.6733967025 | Human_heart | SIMS |
| 0.7359383378 | Human_heart | scNym |
| 0.7595371972 | Human_heart | scNym |
| 0.7527092312 | Human_heart | scNym |
| 0.761362745 | Human_heart | scNym |

| | | |
|---|---|---|
| **0.7376867833** | Human_heart | scNym |
| **0.4755880961** | Human_heart | Scanvi |
| **0.4335355828** | Human_heart | Scanvi |
| **0.4358046662** | Human_heart | Scanvi |
| **0.4826174999** | Human_heart | Scanvi |
| **0.4398545121** | Human_heart | Scanvi |
| **0.7529132178** | Human_heart | SciBet |
| **0.7497644457** | Human_heart | SciBet |
| **0.7465378152** | Human_heart | SciBet |
| **0.7478067598** | Human_heart | SciBet |
| **0.7498167832** | Human_heart | SciBet |
| **0.8358666461** | Human_heart | Seurat |
| **0.8345380698** | Human_heart | Seurat |
| **0.8339608111** | Human_heart | Seurat |
| **0.8353435788** | Human_heart | Seurat |
| **0.8380069703** | Human_heart | Seurat |
| **0.7437973944** | Human_heart | SingleR |
| **0.7413119593** | Human_heart | SingleR |
| **0.7444221778** | Human_heart | SingleR |
| **0.746006217** | Human_heart | SingleR |
| **0.7447123412** | Human_heart | SingleR |
| **0.692** | PBMC_68K | SIMS |
| **0.701** | PBMC_68K | SIMS |
| **0.705** | PBMC_68K | SIMS |
| **0.699** | PBMC_68K | SIMS |
| **0.697** | PBMC_68K | SIMS |
| **0.6887** | PBMC_68K | scBERT |
| **0.69** | PBMC_68K | scBERT |
| **0.6895** | PBMC_68K | scBERT |
| **0.6921** | PBMC_68K | scBERT |
| **0.6935** | PBMC_68K | scBERT |
| **0.6272890391** | PBMC_68K | scNym |
| **0.6272718658** | PBMC_68K | scNym |
| **0.63089981** | PBMC_68K | scNym |
| **0.6238039372** | PBMC_68K | scNym |
| **0.6291427101** | PBMC_68K | scNym |

| | | |
|---|---|---|
| **0.515314189** | PBMC_68K | Scanvi |
| **0.615996898** | PBMC_68K | Scanvi |
| **0.6137325** | PBMC_68K | Scanvi |
| **0.60695337** | PBMC_68K | Scanvi |
| **0.6253614** | PBMC_68K | Scanvi |
| **0.6664182** | PBMC_68K | SciBet |
| **0.6558428** | PBMC_68K | SciBet |
| **0.6558846** | PBMC_68K | SciBet |
| **0.6481533** | PBMC_68K | SciBet |
| **0.6577924** | PBMC_68K | SciBet |
| **0.581276** | PBMC_68K | Seurat |
| **0.5791013** | PBMC_68K | Seurat |
| **0.5732654** | PBMC_68K | Seurat |
| **0.5775451** | PBMC_68K | Seurat |
| **0.5880288** | PBMC_68K | Seurat |
| **0.5509479** | PBMC_68K | SingleR |
| **0.5447428** | PBMC_68K | SingleR |
| **0.5454915** | PBMC_68K | SingleR |
| **0.5526609** | PBMC_68K | SingleR |
| **0.5407837** | PBMC_68K | SingleR |

**Supplemental Table 3. Accuracies by cell type for sample H200.1025 predictions training SIMS on Sample H200.1023.**

| Cell Type | Precision | Recall | F1-score | Support |
|---|---|---|---|---|
| **Astrocyte** | 0.951 | 0.993 | 0.971 | 446 |
| **Endothelial Cell** | 0.917 | 0.550 | 0.688 | 20 |
| **Intratelencephalic Projection Neuron** | 0.980 | 0.925 | 0.952 | 4575 |
| **Layer 4 Intratelencephalic Projection Neuron** | 0.787 | 0.919 | 0.848 | 936 |
| **Layer 5 Extratelencephalic Projection Neuron** | 0.724 | 0.677 | 0.700 | 31 |
| **Layers 5/6 CAR3+ Intratelencephalic Projection Neuron** | 0.969 | 0.916 | 0.942 | 239 |
| **Layers 5/6 Near Projecting Neuron** | 0.949 | 0.931 | 0.940 | 160 |
| **Layer 6 Corticothalamic Projection Neuron** | 0.961 | 0.914 | 0.937 | 627 |
| **Layer 6b Projection Neuron** | 0.904 | 0.943 | 0.923 | 229 |
| **LAMP5+ Interneuron** | 0.920 | 0.987 | 0.953 | 549 |
| **Microglia** | 0.941 | 0.989 | 0.965 | 179 |
| **Oligodendrocyte Precursor Cell** | 0.957 | 0.975 | 0.966 | 204 |
| **Oligodendrocyte** | 0.894 | 0.981 | 0.935 | 463 |
| **PAX6+ Interneuron** | 0.810 | 0.914 | 0.859 | 70 |
| **PVALB+ Interneuron** | 0.955 | 0.969 | 0.962 | 618 |
| **Pericyte** | 0.500 | 0.500 | 0.500 | 4 |
| **SST+ Interneuron** | 0.950 | 0.982 | 0.966 | 614 |
| **VIP+ Interneuron** | 0.956 | 0.943 | 0.949 | 915 |
| **Vascular Leptomeningeal Cell** | 0.000 | 0.000 | 0.000 | 1 |
| **Accuracy** | 0.940 | 0.940 | 0.940 | 0.9399816176 |
| **Macro Avg** | 0.843 | 0.843 | 0.840 | 10880 |
| **Weighted Avg** | 0.944 | 0.940 | 0.941 | 10880 |

**Supplemental Table 4. Accuracies by cell type for sample H200.1030 predictions training SIMS on Sample H200.1023.**

| Cell Type | Precision | Recall | F1-score | Support |
|---|---|---|---|---|
| Astrocyte | 0.935 | 0.950 | 0.943 | 458 |
| Endothelial Cell | 0.970 | 0.865 | 0.914 | 37 |
| Intratelencephalic Projection Neuron | 0.970 | 0.953 | 0.962 | 8635 |
| Layer 4 Intratelencephalic Projection Neuron | 0.837 | 0.867 | 0.852 | 1658 |
| Layer 5 Extratelencephalic Projection Neuron | 0.860 | 0.557 | 0.676 | 88 |
| Layers 5/6 CAR3+ Intratelencephalic Projection Neuron | 0.970 | 0.925 | 0.947 | 425 |
| Layers 5/6 Near Projecting Neuron | 0.967 | 0.962 | 0.965 | 340 |
| Layer 6 Corticothalamic Projection Neuron | 0.953 | 0.933 | 0.943 | 1018 |
| Layer 6b Projection Neuron | 0.863 | 0.945 | 0.902 | 326 |
| LAMP5+ Interneuron | 0.949 | 0.969 | 0.959 | 991 |
| Microglia | 0.959 | 0.922 | 0.940 | 307 |
| Oligodendrocyte Precursor Cell | 0.944 | 0.951 | 0.947 | 264 |
| Oligodendrocyte | 0.909 | 0.973 | 0.940 | 736 |
| PAX6+ Interneuron | 0.848 | 0.803 | 0.825 | 132 |
| PVALB+ Interneuron | 0.954 | 0.971 | 0.962 | 1119 |
| Pericyte | 0.813 | 0.929 | 0.867 | 14 |
| SST+ Interneuron | 0.933 | 0.961 | 0.946 | 938 |
| VIP+ Interneuron | 0.945 | 0.963 | 0.954 | 1320 |
| Vascular Leptomeningial Cell | 0.000 | 0.000 | 0.000 | 0 |
| Accuracy | 0.944 | 0.944 | 0.944 | 0.9437413591 |
| Macro Avg | 0.873 | 0.863 | 0.865 | 18806 |
| Weighted Avg | 0.945 | 0.944 | 0.944 | 18806 |

**Supplemental Table 5. Accuracies by cell type for sample H200.1023 predictions training SIMS on Sample H200.1025.**

| Cell Type | Precision | Recall | F1-score | Support |
|---|---|---|---|---|
| **Astrocyte** | 0.941 | 0.828 | 0.881 | 291.000 |
| **Endothelial Cell** | 0.680 | 0.944 | 0.791 | 18.000 |
| **Intratelencephalic Projection Neuron** | 0.962 | 0.959 | 0.961 | 9066.000 |
| **Layer 4 Intratelencephalic Projection Neuron** | 0.724 | 0.774 | 0.748 | 857.000 |
| **Layer 5 Extratelencephalic Projection Neuron** | 0.306 | 0.595 | 0.404 | 37.000 |
| **Layers 5/6 CAR3+ Intratelencephalic Projection Neuron** | 0.964 | 0.983 | 0.974 | 414.000 |
| **Layers 5/6 Near Projecting Neuron** | 0.888 | 0.846 | 0.866 | 337.000 |
| **Layer 6 Corticothalamic Projection Neuron** | 0.962 | 0.882 | 0.920 | 1050.000 |
| **Layer 6b Projection Neuron** | 0.818 | 0.868 | 0.843 | 456.000 |
| **LAMP5+ Interneuron** | 0.941 | 0.877 | 0.908 | 894.000 |
| **Microglia** | 0.944 | 0.981 | 0.962 | 257.000 |
| **Oligodendrocyte Precursor Cell** | 0.900 | 0.978 | 0.937 | 275.000 |
| **Oligodendrocyte** | 0.921 | 0.956 | 0.938 | 611.000 |
| **PAX6+ Interneuron** | 0.603 | 0.689 | 0.643 | 106.000 |
| **PVALB+ Interneuron** | 0.965 | 0.945 | 0.955 | 1057.000 |
| **Pericyte** | 0.000 | 0.000 | 0.000 | 3.000 |
| **SST+ Interneuron** | 0.937 | 0.949 | 0.943 | 747.000 |
| **VIP+ Interneuron** | 0.932 | 0.947 | 0.940 | 1265.000 |
| **Vascular Leptomeningial Cell** | 0.000 | 0.000 | 0.000 | 5.000 |
| **Accuracy** | 0.931 | 0.931 | 0.931 | 0.931 |
| **Macro Avg** | 0.757 | 0.790 | 0.769 | 17746.000 |
| **Weighted Avg** | 0.934 | 0.931 | 0.932 | 17746.000 |

**Supplemental Table 6. Accuracies by cell type for sample H200.1030 predictions training SIMS on Sample H200.1025.**

| Cell Type | Precision | Recall | F1-score | Support |
|---|---|---|---|---|
| **Astrocyte** | 0.929 | 0.887 | 0.908 | 487.000 |
| **Endothelial Cell** | 0.879 | 0.853 | 0.866 | 34.000 |
| **Intratelencephalic Projection Neuron** | 0.963 | 0.955 | 0.959 | 8562.000 |
| **Layer 4 Intratelencephalic Projection Neuron** | 0.822 | 0.912 | 0.865 | 1546.000 |
| **Layer 5 Extratelencephalic Projection Neuron** | 0.509 | 0.580 | 0.542 | 50.000 |
| **Layers 5/6 CAR3+ Intratelencephalic Projection Neuron** | 0.943 | 0.992 | 0.967 | 385.000 |
| **Layers 5/6 Near Projecting Neuron** | 0.882 | 0.782 | 0.829 | 381.000 |
| **Layer 6 Corticothalamic Projection Neuron** | 0.977 | 0.870 | 0.921 | 1119.000 |
| **Layer 6b Projection Neuron** | 0.711 | 0.870 | 0.783 | 292.000 |
| **LAMP5+ Interneuron** | 0.941 | 0.889 | 0.914 | 1071.000 |
| **Microglia** | 0.966 | 0.990 | 0.978 | 288.000 |
| **Oligodendrocyte Precursor Cell** | 0.876 | 0.925 | 0.900 | 252.000 |
| **Oligodendrocyte** | 0.933 | 0.963 | 0.948 | 763.000 |
| **PAX6+ Interneuron** | 0.576 | 0.610 | 0.593 | 118.000 |
| **PVALB+ Interneuron** | 0.953 | 0.941 | 0.947 | 1153.000 |
| **Pericyte** | 0.000 | 0.000 | 0.000 | 5.000 |
| **SST+ Interneuron** | 0.946 | 0.934 | 0.940 | 979.000 |
| **VIP+ Interneuron** | 0.929 | 0.949 | 0.939 | 1316.000 |
| **Vascular Leptomeningial Cell** | 0.000 | 0.000 | 0.000 | 5.000 |
| **Accuracy** | 0.931 | 0.931 | 0.931 | 0.931 |
| **Macro Avg** | 0.776 | 0.784 | 0.779 | 18806.000 |
| **Weighted Avg** | 0.934 | 0.931 | 0.932 | 18806.000 |

**Supplemental Table 7. Accuracies by cell type for sample H200.1023 predictions training SIMS on Sample H200.1030.**

| Cell Type | Precision | Recall | F1-score | Support |
|---|---|---|---|---|
| **Astrocyte** | 0.969 | 0.961 | 0.965 | 258 |
| **Endothelial Cell** | 0.800 | 0.769 | 0.784 | 26 |
| **Intratelencephalic Projection Neuron** | 0.985 | 0.961 | 0.973 | 9264 |
| **Layer 4 Intratelencephalic Projection Neuron** | 0.713 | 0.870 | 0.783 | 751 |
| **Layer 5 Extratelencephalic Projection Neuron** | 0.764 | 0.873 | 0.815 | 63 |
| **Layers 5/6 CAR3+ Intratelencephalic Projection Neuron** | 0.967 | 0.938 | 0.952 | 435 |
| **Layers 5/6 Near Projecting Neuron** | 0.963 | 0.978 | 0.970 | 316 |
| **Layer 6 Corticothalamic Projection Neuron** | 0.953 | 0.947 | 0.950 | 969 |
| **Layer 6b Projection Neuron** | 0.919 | 0.939 | 0.929 | 474 |
| **LAMP5+ Interneuron** | 0.975 | 0.988 | 0.981 | 822 |
| **Microglia** | 0.940 | 0.980 | 0.960 | 256 |
| **Oligodendrocyte Precursor Cell** | 0.943 | 0.949 | 0.946 | 297 |
| **Oligodendrocyte** | 0.946 | 0.982 | 0.964 | 611 |
| **PAX6+ Interneuron** | 0.975 | 0.781 | 0.868 | 151 |
| **PVALB+ Interneuron** | 0.971 | 0.970 | 0.971 | 1036 |
| **Pericyte** | 0.667 | 0.615 | 0.640 | 13 |
| **SST+ Interneuron** | 0.963 | 0.967 | 0.965 | 754 |
| **VIP+ Interneuron** | 0.965 | 0.993 | 0.979 | 1249 |
| **Vascular Leptomeningial Cell** | 0.000 | 0.000 | 0.000 | 1 |
| **Accuracy** | 0.958 | 0.958 | 0.958 | 0.9582441113 |
| **Macro Avg** | 0.862 | 0.866 | 0.863 | 17746 |
| **Weighted Avg** | 0.961 | 0.958 | 0.959 | 17746 |

**Supplemental Table 8. Accuracies by cell type for sample H200.1025 predictions training SIMS on Sample H200.1030.**

| Cell Type | Precision | Recall | F1-score | Support |
|---|---|---|---|---|
| Astrocyte | 0.942 | 0.989 | 0.965 | 444 |
| Endothelial Cell | 1.000 | 0.923 | 0.960 | 13 |
| Intratelencephalic Projection Neuron | 0.979 | 0.934 | 0.956 | 4526 |
| Layer 4 Intratelencephalic Projection Neuron | 0.835 | 0.908 | 0.870 | 1005 |
| Layer 5 Extratelencephalic Projection Neuron | 0.793 | 0.852 | 0.821 | 27 |
| Layers 5/6 CAR3+ Intratelencephalic Projection Neuron | 0.947 | 0.951 | 0.949 | 225 |
| Layers 5/6 Near Projecting Neuron | 0.962 | 0.987 | 0.974 | 153 |
| Layer 6 Corticothalamic Projection Neuron | 0.958 | 0.953 | 0.956 | 599 |
| Layer 6b Projection Neuron | 0.895 | 0.934 | 0.915 | 229 |
| LAMP5+ Interneuron | 0.944 | 0.977 | 0.960 | 569 |
| Microglia | 0.920 | 0.966 | 0.943 | 179 |
| Oligodendrocyte Precursor Cell | 0.947 | 0.980 | 0.963 | 201 |
| Oligodendrocyte | 0.939 | 0.984 | 0.961 | 485 |
| PAX6+ Interneuron | 0.899 | 0.789 | 0.840 | 90 |
| PVALB+ Interneuron | 0.955 | 0.955 | 0.955 | 627 |
| Pericyte | 0.750 | 0.500 | 0.600 | 6 |
| SST+ Interneuron | 0.970 | 0.976 | 0.973 | 631 |
| VIP+ Interneuron | 0.952 | 0.987 | 0.970 | 871 |
| Vascular Leptomeningial Cell | 0.000 | 0.000 | 0.000 | 0 |
| Accuracy | 0.948 | 0.948 | 0.948 | 0.9481617647 |
| Macro Avg | 0.873 | 0.871 | 0.870 | 10880 |
| Weighted Avg | 0.950 | 0.948 | 0.949 | 10880 |

### Supplemental Table 9. Accuracies by cell type for E18 and P1 samples predictions training SIMS on E18 and P1 samples

| Cell Type | Precision | Recall | F1-score | Support |
|---|---|---|---|---|
| **Astrocytes** | 0.920 | 0.837 | 0.876 | 754.000 |
| **Corticothalamic Projection Neurons** | 0.930 | 0.871 | 0.900 | 519.000 |
| **Cajal Retzius Cells** | 0.984 | 0.984 | 0.984 | 64.000 |
| **Cycling Glial Cells** | 0.593 | 0.596 | 0.594 | 235.000 |
| **Deep Layer Callosal Projection Neurons** | 0.908 | 0.692 | 0.785 | 412.000 |
| **Endothelial cells** | 1.000 | 1.000 | 1.000 | 24.000 |
| **Ependymocytes** | 0.000 | 0.000 | 0.000 | 26.000 |
| **Intermediate Progenitors** | 0.422 | 0.595 | 0.493 | 190.000 |
| **Interneurons** | 0.937 | 0.971 | 0.954 | 2498.000 |
| **Layer 4 Neurons** | 0.795 | 0.572 | 0.665 | 1325.000 |
| **Layer 6b Projection Neurons** | 0.892 | 0.733 | 0.805 | 45.000 |
| **Microglia** | 0.975 | 0.975 | 0.975 | 40.000 |
| **Migrating Neurons** | 0.653 | 0.809 | 0.723 | 918.000 |
| **Near Projecting Neurons** | 0.726 | 0.794 | 0.759 | 180.000 |
| **Oligodendrocytes** | 0.927 | 0.948 | 0.937 | 213.000 |
| **Pericytes** | 0.966 | 1.000 | 0.982 | 28.000 |
| **Subcerebral Projection Neurons** | 0.944 | 0.941 | 0.942 | 304.000 |
| **Upper Layers Callosal Projection Neurons** | 0.845 | 0.882 | 0.863 | 4427.000 |
| **Accuracy** | 0.842 | 0.842 | 0.842 | 0.842 |
| **Macro Avg** | 0.801 | 0.789 | 0.791 | 12202.000 |
| **Weighted Avg** | 0.845 | 0.842 | 0.840 | 12202.000 |

**Supplemental Table 10. Accuracies by cell type for E18 samples predictions training SIMS on P1 samples.**

| Cell Type | Precision | Recall | F1-score | Support |
|---|---|---|---|---|
| **Astrocytes** | 0.309 | 0.927 | 0.463 | 41.000 |
| **Corticothalamic Projection Neurons** | 0.787 | 0.799 | 0.793 | 422.000 |
| **Cajal Retzius Cells** | 0.927 | 0.950 | 0.938 | 40.000 |
| **Cycling Glial Cells** | 0.238 | 0.242 | 0.240 | 62.000 |
| **Deep Layer Callosal Projection Neurons** | 0.843 | 0.566 | 0.677 | 343.000 |
| **Endothelial cells** | 0.941 | 1.000 | 0.970 | 16.000 |
| **Intermediate Progenitors** | 0.583 | 0.082 | 0.144 | 85.000 |
| **Interneurons** | 0.928 | 0.931 | 0.929 | 1209.000 |
| **Layer 4 Neurons** | 0.781 | 0.243 | 0.371 | 864.000 |
| **Layer 6b Projection Neurons** | 0.935 | 0.707 | 0.806 | 41.000 |
| **Microglia** | 1.000 | 0.952 | 0.976 | 21.000 |
| **Migrating Neurons** | 0.263 | 0.921 | 0.410 | 202.000 |
| **Near Projecting Neurons** | 0.365 | 0.182 | 0.243 | 148.000 |
| **Oligodendrocytes** | 0.875 | 0.850 | 0.863 | 107.000 |
| **Pericytes** | 1.000 | 0.867 | 0.929 | 15.000 |
| **Subcerebral Projection Neurons** | 0.976 | 0.864 | 0.917 | 286.000 |
| **Upper Layers Callosal Projection Neurons** | 0.752 | 0.827 | 0.788 | 3086.000 |
| **Accuracy** | 0.736 | 0.736 | 0.736 | 0.736 |
| **Macro Avg** | 0.736 | 0.701 | 0.674 | 6988.000 |
| **Weighted Avg** | 0.776 | 0.736 | 0.727 | 6988.000 |

**Supplemental Table 11. Accuracies by cell type for GM8330-derived organoids predictions training SIMS on 11A-derived organoids.**

| Classification Report | precision | recall | f1-score | support |
|---|---|---|---|---|
| Cycling | 0.63 | 0.68 | 0.65 | 864 |
| Intermediate Progenitor Cells | 0.56 | 0.66 | 0.61 | 679 |
| Immature Callosal Projection Neurons | 0.42 | 0.89 | 0.57 | 629 |
| Immature Interneurons | 0.83 | 0.90 | 0.86 | 4299 |
| Immature Projection Neurons | 0.12 | 0.33 | 0.17 | 465 |
| Radial Glia | 0.49 | 0.09 | 0.15 | 2647 |
| Ventral Precursors | 0.80 | 0.65 | 0.72 | 1104 |
| Outer Radial Glia/Astroglia | 0.79 | 0.85 | 0.82 | 3534 |
| accuracy | 0.67 | 14221 | | |
| macro avg | 0.58 | 0.63 | 0.57 | 14221 |
| weighted avg | 0.69 | 0.67 | 0.65 | 14221 |

**Supplemental Table 12. Accuracies by cell type for PGP1-derived organoids predictions training SIMS on 11A-derived organoids.**

| Classification Report | precision | recall | f1-score | support |
|---|---|---|---|---|
| Cycling | 0.89 | 0.63 | 0.74 | 1750 |
| Intermediate Progenitor Cells | 0.12 | 0.91 | 0.22 | 170 |
| Immature Callosal Projection Neurons | 0.86 | 0.76 | 0.81 | 4144 |
| Immature Interneurons | 0.55 | 0.98 | 0.70 | 2261 |
| Immature Projection Neurons | 0.48 | 0.51 | 0.49 | 1362 |
| Radial Glia | 0.01 | 0.02 | 0.01 | 418 |
| Ventral Precursors | 0.77 | 0.70 | 0.73 | 1329 |
| Outer Radial Glia/Astroglia | 0.97 | 0.70 | 0.82 | 9171 |
| accuracy | 0.71 | 20605 | | |
| macro avg | 0.58 | 0.65 | 0.56 | 20605 |
| weighted avg | 0.82 | 0.71 | 0.75 | 20605 |

**Supplemental Table 13. Accuracies by cell type for GM8330-derived organoids predictions training SIMS on 11A-derived organoids after reclassification of 11A cells.**

| Classification Report | precision | recall | f1-score | support |
|---|---|---|---|---|
| Cycling | 0.64 | 0.64 | 0.64 | 864 |
| Intermediate Progenitor Cells | 0.57 | 0.69 | 0.62 | 679 |
| Immature Callosal Projection Neurons | 0.39 | 0.83 | 0.53 | 629 |
| Immature Interneurons | 0.86 | 0.88 | 0.87 | 4504 |
| Immature Projection Neurons | 0.11 | 0.39 | 0.17 | 163 |
| Radial Glia | 0.83 | 0.33 | 0.47 | 2744 |
| Ventral Precursors | 0.78 | 0.71 | 0.75 | 1104 |
| Outer Radial Glia/Astroglia | 0.77 | 0.86 | 0.81 | 3534 |
| accuracy | 0.72 | 14221 | | |
| macro avg | 0.62 | 0.66 | 0.61 | 14221 |
| weighted avg | 0.77 | 0.72 | 0.72 | 14221 |

**Supplemental Table 14. Accuracies by cell type for PGP1-derived organoids predictions training SIMS on 11A-derived organoids**

| Classification Report | precision | recall | f1-score | support |
|---|---|---|---|---|
| Cycling | 0.91 | 0.61 | 0.73 | 1750 |
| Intermediate Progenitor Cells | 0.14 | 0.93 | 0.24 | 170 |
| Immature Callosal Projection Neurons | 0.87 | 0.82 | 0.84 | 4144 |
| Immature Interneurons | 0.58 | 0.98 | 0.73 | 2261 |
| Immature Projection Neurons | 0.63 | 0.39 | 0.49 | 1362 |
| Radial Glia | 0.17 | 0.61 | 0.27 | 418 |
| Ventral Precursors | 0.74 | 0.79 | 0.76 | 1329 |
| Outer Radial Glia/Astroglia | 0.97 | 0.73 | 0.83 | 9171 |
| accuracy | 0.74 | 20605 | | |
| macro avg | 0.63 | 0.73 | 0.61 | 20605 |
| weighted avg | 0.84 | 0.74 | 0.77 | 20605 |
